# HIV-1 and microglia: EcoHIV and HIV-1 transgenic rats

**DOI:** 10.1101/2020.11.03.365494

**Authors:** Hailong Li, Kristen A. McLaurin, Jessica M. Illenberger, Charles F. Mactutus, Rosemarie M. Booze

## Abstract

The persistence of HIV-1 viral reservoirs in the brain, despite treatment with combination antiretroviral therapy (cART), remains a critical roadblock for the development of a novel cure strategy for HIV-1. To enhance our understanding of viral reservoirs, two complementary studies were conducted to 1) evaluate the HIV-1 mRNA neuroanatomical distribution pattern and major cell type expressing HIV-1 mRNA in the HIV-1 transgenic (Tg) rat (i.e., under conditions of latent infection), and 2) to validate our findings by developing and critically testing a novel biological system to model active HIV-1 infection in the rat. First, a restricted, region-specific HIV-1 mRNA distribution pattern was observed in the HIV-1 Tg rat. Microglia were the predominant cell type expressing HIV-1 mRNA in the HIV-1 Tg rat. Second, we developed and critically tested a novel biological system to model key aspects of HIV-1 by infusing F344/N control rats with chimeric HIV (EcoHIV). *In vitro,* primary cultured microglia were treated with EcoHIV revealing prominent expression within 24 hours of infection. *In vivo,* EcoHIV expression was observed seven days after stereotaxic injections. Following EcoHIV infection, microglia were the major cell type expressing HIV-1 mRNA, results which are consistent with observations in the HIV-1 Tg rat. Within eight weeks of infection, EcoHIV rats exhibited neurocognitive impairments, synaptic dysfunction, which may result from activation of the NogoA-NgR3/PirB-RhoA signaling pathway, and neuroinflammation. Collectively, these studies enhance our understanding of HIV-1 viral reservoirs in the brain and offer a novel biological system to model HIV-associated neurocognitive disorders and associated comorbidities (i.e., drug abuse) in rats.

## INTRODUCTION

Human immunodeficiency virus type 1 (HIV-1), which afflicts approximately 37.9 million individuals worldwide (UNAIDS, 2019), continues to be a public health crisis. Within three to five days of HIV-1 infection^1,2^, infected CD4+ cells and monocytes transmigrate across the blood-brain barrier^3^ infecting astrocytes, perivascular macrophages and microglial cells^4^. Although combination antiretroviral therapy (cART), the primary treatment regimen for HIV-1, suppresses viral replication in the periphery, it fails to penetrate the blood-brain barrier^5^. Consequently, the central nervous system (CNS) acts as a latent viral reservoir for HIV-1^6^; a factor that is associated with the emergence of virus resistance^7,8^, as well as the development and progression of HIV-1 associated neurocognitive disorders (HAND)^9,10^.

Microglia cells are innate immune cells in the CNS that represent 5-20% of adult brain cells^11^. Key structural characteristics of microglia, including a long half-life^12^, ability to undergo cell division^13^ and susceptibility to HIV-1 infection^14^, support microglia as one of the latent viral reservoirs for HIV-1 in the brain^15^. Specifically, utilization of a highly sensitive *in situ* hybridization technique, in combination with immunohistochemistry, revealed that brain macrophages and microglia, but not astrocytes, were harboring HIV-1 DNA in the brain^16^; results which are consistent with earlier findings in HIV-1 seropositive individuals with HIV-1 encephalitis^4,17^. Furthermore, aberrant microglial activation is associated with alterations in synaptic function^18^ neurotransmitter excitotoxicity^19^, and neuroinflammation^20^; functional alterations which have also been observed in HIV-1 seropositive individuals^21,22,23^. Altogether, microglia exhibit structural and functional characteristics which meet the criteria of a latent viral reservoir for HIV-1 infection in the brain^24^.

Studying CNS infection of HIV-1 in humans, however, is limited by ethical and technical constraints, supporting the importance of biological systems to model key aspects of HIV-1 and HAND. Specifically, biological systems, including macaques infected with Simian immunodeficiency virus^25,26,27^, humanized mice^28^, and EcoHIV infected mice^29^ have provided strong evidence for the infection of microglial cells and development of a viral reservoir in HIV-1. However, to date, no study has systematically evaluated the major cell type expressing HIV-1 mRNA in the HIV-1 transgenic (Tg) rat, which has been supported as a biological system to model HIV-1 associated neurocognitive disorders (HAND) in the post-cART era ^30,31,32,33^. In light of these gaps in our knowledge, two complementary experiments were conducted to 1) evaluate the HIV-1 mRNA neuroanatomical distribution pattern and major cell type expressing HIV-1 mRNA in the HIV-1 transgenic (Tg) rat, and 2) to validate our findings by developing and critically testing a novel model of HIV-1 infection in the rat (i.e., chimeric HIV (EcoHIV)).

First, the HIV-1 Tg rat, a biological system to model latent infection, was utilized to characterize the brain regions, and major cell type, expressing HIV-1 mRNA. The HIV-1 Tg rat, originally reported by Reid et al.^34^, resembles HIV-1 seropositive individuals on cART and has been utilized to investigate neurocognitive^30,31,33^ and neuroanatomical^32,35, 36^ alterations associated with HAND. The functional deletion of *gag*- and *pol*-, a reverse transcriptase, precludes viral replication rendering the HIV-1 Tg rat noninfectious. HIV-1 viral transcripts have been observed throughout the body of the HIV-1 Tg rat, including in the spleen, liver, thymus, lymph nodes, and kidneys^34, 37,38^. Furthermore, viral proteins have also been detected in the tissues, macrophages, and in serum of these animals^37,39^. Although the HIV-1 provirus is present in all cells, the actual HIV-1 mRNA expression pattern in the brain of the HIV-1 Tg rat remains unclear; a need which was addressed in the present study by utilizing RNA *in situ* hybridization (RNAscope), a highly sensitive and innovative technique.

Second, we developed a novel biological system to model active HIV-1 infection in the rat using chimeric HIV (EcoHIV). EcoHIV infection in mice, originally reported by Potash et al.^40^, recapitulates many of the clinical features of HIV-1 commonly observed in humans, including lymphocyte and macrophage infection, induction of antiviral immune responses, neuro-invasiveness, and an increase of inflammatory and antiviral factors in brain^29,41^. Despite the significant utility of EcoHIV mice, extending this biological system to rats, which are more commonly utilized for studies of HAND and drug abuse, would be advantageous. Thus, the present study utilized *in vitro* and *in vivo* techniques to critically test and characterize EcoHIV infection in rats. Understanding HIV-1 viral reservoirs in the brain and the development of a novel biological system to assess HAND and associated comorbidities (i.e., drug abuse) may afford innovative targets for the development of therapeutics and/or cure strategies.

## METHODS

### Animals

All rats in the current experiments were pair-housed in AAALAC-accredited facilities using guidelines established by the National Institute of Health. The environmental conditions were targeted at a 12h:12h light/dark cycle with lights on at 700 h (EST), 21°± 2°C, and 50% ± 10% relative humidity. Animals had *ad libitum* access to rodent chow (2020X (Harlan Teklad, Madison, WI)) and water. Fischer (F344/N) and HIV-1 Tg rats were obtained from Envigo Laboratories (Indianapolis, IN). The project protocol was approved by the Institutional Animal Care and Use Committee (IACUC) at the University of South Carolina (federal assurance number: # D16-00028).

### Experimental Design

A schematic of the experimental design is presented in Figure 1.

**Figure 1.**
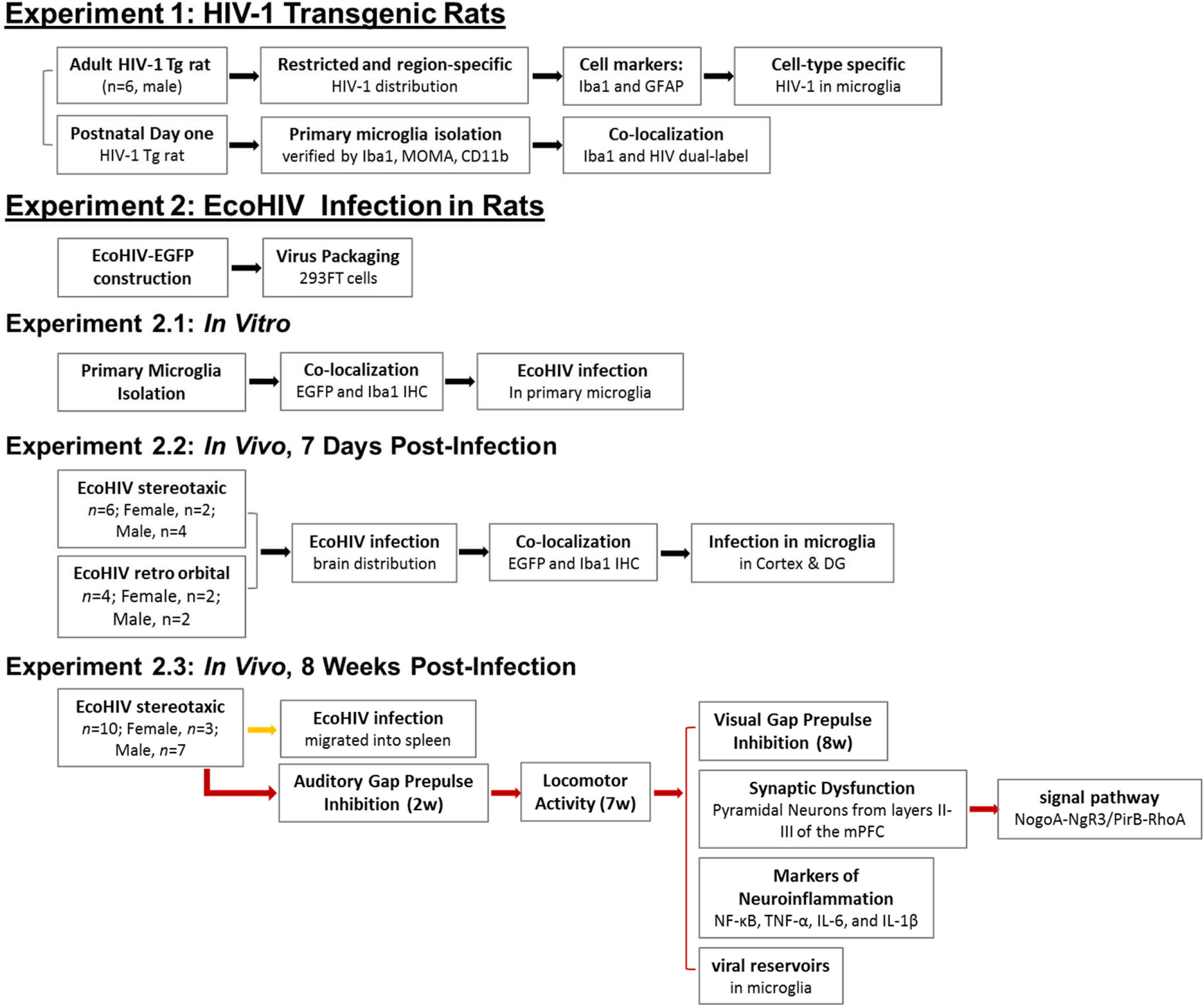
Schematic of the experimental design.

### Experiment 1

#### RNAscope In Situ Hybridization

RNAscope *in situ* hybridization was used to compare the expression of HIV-1 mRNA across different brain regions of HIV-1 Tg animals. The RNAscope *in situ* hybridization protocol was described in detail by Li et al.^42^.

In brief, HIV- 1 Tg rats (*n*=6) were deeply anesthetized using 5% sevoflurane and sacrificed. The brain tissue was removed and frozen in liquid nitrogen within 5 minutes of sacrifice. 30 μm sagittal sections were cut using a cryostat and mounted onto SuperFrost Plus slides, which were dried at −20°C for 10 minutes. Subsequently, slides were immersed in 4% paraformaldehyde for 1 hour at 4°C followed by an increasing ethanol gradient (50%, 70%, 100% EtOH). Next, a pretreatment reagent, (i.e., Protease IV Reagent, RNAscope Fluorescence Multiplex Kit, Advanced Cell Diagnostics, Newark, CA) was applied to sections. Sections were incubated for 30 minutes at room temperature and then hybridized with specific probes for HIV-1 viral proteins (see Supplementary Table S2). Slides were placed in the HybEZ Oven (Advanced Cell Diagnostics, Newark, CA) and incubated at 40°C for 2 hours. Subsequently, signals were amplified using “Amp 1-FL”, “Amp 2-FL”, “Amp 3-FL”, and “Amp 4-FL-Alt A” reagents provided in the RNAscope Fluorescence Multiplex Kit, were applied to each section and incubated for either 30 minutes (“Amp 1-FL”, “Amp 3-FL”) or 15 minutes (“Amp 2-FL”, “Amp 4-FL-Alt A”) at 40°C. Slides were mounted with Pro-Long Gold Antifade (Invitrogen, Carlsbad, CA), coverslipped and stored in the dark at 4°C until dry. Z-stack images were obtained using a 60× objective on a Nikon TE-2000E confocal microscope utilizing Nikon's EZ-C1 software (version 3.81b).

The primary cell type expressing HIV-1 mRNA in the HIV-1 Tg rat was examined using two methods. First, RNAscope *in situ* hybridization images were compared to those of adjacent sections, which were processed with immunohistochemical staining for microglia (Iba1) and astrocytes (GFAP; methodology detailed below); a method utilized due to technological limitations. Second, RNAscope dual-labelling was utilized to more accurately assess co-localization of expression for HIV-1 mRNA and specific cell type markers (i.e., microglia: Iba1 mRNA, astrocytes: GFAP mRNA) in primary microglia cultures from HIV-1 Tg rats (methodology detailed below) and brain tissue from HIV-1 Tg rats.

#### Immunohistochemical Staining

In addition to RNAscope *in situ* hybridization, immunohistochemical staining was used to examine the localization of HIV-1 mRNA within different cell types (i.e., microglia, astrocytes) in brain tissue from the HIV-1 Tg rat. After perfusion (*n*=6 male, HIV-1 Tg animals), brains were removed and post fixed overnight in 4% chilled paraformaldehyde. Brain tissue was sectioned into 50μm thick coronal slices using a vibratome (PELCO easiSlicer™, *TED PELLA, INC.*). Brain sections were incubated overnight at 4°C with rabbit anti-Iba1 (ab178847, Abcam, Cambridge, United Kingdom) and rabbit anti-GFAP (ab7260, Abcam, Cambridge, United Kingdom). Subsequently, Alexa Fluor 594 goat anti-rabbit IgG (A11012, Invitrogen) was applied to brain sections and incubated for 4 hours at room temperature. Z-stack images were obtained using a 60× objective on a Nikon TE-2000E confocal microscope utilizing Nikon’s EZ-C1 software (version 3.81b).

#### Primary Microglia Isolation and Purification

To validate our observations, microglia cultures were harvested from HIV-1 Tg rats to examine the expression of HIV-1 mRNA.

First, mixed glia were isolated from the HIV-1 Tg rat on postnatal day (PD) 1, using a protocol that was modified from Moussaud, *et al*.^43^. In brief, after the rats were deeply anesthetized, the brain was aseptically removed and the meninges were gently peeled off. A piece of the cortex was transferred to a small Petri dish containing 2 ml of DMEM/F12 medium, finely minced with a blade, transferred into a 15 mL centrifuge tube with DMEM/F12 medium and 0.5% Trypsin/EDTA, and incubated at 37°C in a humidified 5% CO_2_ atmosphere for 20 minutes. The single cell suspension was gently agitated by moving the solution up and down 20 times using a 3.5 ml plastic transfer pipette and centrifuged at 1500 rpm for 4 minutes. The cell pellet was resuspended in DMEM/F12 medium with 10% FBS and the cells were seeded into a flask coated with poly-L-lysine. Cells were incubated overnight at 37°C in a humidified 5% CO_2_ atmosphere. After incubating overnight, the DMEM/F12 medium with 10% FBS was replaced. Subsequently, the DMEM/F12 medium with 10% FBS was changed every three days.

Once the mixed glia reached 90% confluency (after 10-14 days), microglia were detached from the flask using an orbital shaker at 220 rpm for 1 hour (37 °C). The DMEM/F12 medium with 10% FBS was changed no more than 72 hours before isolation. The culture medium was aspirated and centrifuged at 1500 rpm for 4 min. Cells were resuspended and seeded into a cell culture place coated with poly-L-lysine. The purified primary microglia cells were maintained in a humidified 5% CO_2_ atmosphere at 37 °C for 72 hours before experimentation.

Isolation of primary microglia was verified by the immunohistochemical techniques described above (Antibodies: rabbit anti-Iba1, MOMA, and CD11b). Co-localization of HIV-1 viral proteins with microglia was examined using the RNAscope *in situ* hybridization technique described above. Z-stack images were obtained with a Nikon TE-2000E confocal microscope utilizing Nikon’s EZ-C1 software (version 3.81b).

### Experiment 2

#### EcoHIV Virus Construction and Preparation

The lentivirus of EcoHIV-NL4-3-EGFP, generously gifted from Dr. Potash of Icahn School of Medicine at Mount Sinai, was constructed from pNL4-3 in which the fragment of NL4-3 at nucleotide 6310 and NCA-WT at nucleotide 6229 was ligated to generate the chimeric virus. The virus stocks were prepared from the conditional medium after transfection of plasmid DNA into 293FT cells (Lipofectamine™ 3000, Cat. No. L3000015, Invitrogen), and then concentrated using the lenti-x concentrator (Cat. No. 631231, Clontech Laboratories, Mountain View, CA).

#### Experiment 2.1: In Vitro

Primary microglia were isolated and purified using the protocol presented above. Following purification, primary microglia were infected with conditional medium of EcoHIV-EGFP from 293FT cell transfection. The co-localization of EcoHIV infection with microglia was examined using EGFP, which was integral to the lentivirus, and immunohistochemical techniques. Specifically, immunohistochemistry was utilized to visualize microglia using the rabbit anti-Iba1antibody.

#### Experiment 2.2: In Vivo, 7 Days Post-Infection

##### EcoHIV-EGFP virus stereotaxic surgeries

The goal of conducting EcoHIV-EGFP virus stereotaxic surgeries was twofold: 1) To determine whether EcoHIV successfully infects rat brain tissue and 2) To determine the whether EcoHIV expression in the brain is region and/or cell-type specific. Six adult Fisher 344/N rats were stereotaxically injected with the EcoHIV lentivirus (Male, *n*=4; Female, *n*=2). Specifically, animals were anesthetized with sevoflurane (Abbot Laboratories, North Chicago, IL: catalog #035189) and placed in the stereotaxic apparatus (Kopf Instruments, Tujunga, CA: Model 900) with the scalp exposed. Two small 0.40 mm diameter holes were drilled into the skull in neuroanatomical locations relative to Bregma to infuse the viral vectors (1.5 μL EcoHIV NL4-3-EGFP, (1.04 × 10^6^ TU/mL) into the cortex (0.8 mm lateral, 1.2 mm rostral to Bregma, 2.5 mm depth). Viral vectors were infused at a rate of 0.2 μL/min using a 10 μL Hamilton syringe (catalog #1701).

##### RNAscope In Situ Hybridization

Methodology described above for RNAscope dual-labeling was utilized to examine the predominant cell type expressing HIV-1 mRNA seven days after EcoHIV was stereotaxically injected into the brain of F344/N rat.

#### Experiment 2.3: In Vivo, 8 Weeks Post-Infection

After confirming EcoHIV infection in the rat brain, a separate set of animals were used to assess neurocognitive dysfunction and the effect of EcoHIV infection on synaptic dysfunction, signaling pathways, and neuroinflammation. Twenty adult Fischer 344/N rats (Male, *n*=14; Female, *n=*6) were randomly assigned to receive stereotaxic injections of either EcoHIV (*n*=10; Female, *n*=3; Male, *n*=7) or saline (i.e., control; *n*=10; Female, *n*=3; Male, *n*=7).

##### Neurocognitive Assessments: Prepulse Inhibition

Prepulse inhibition (PPI) of the auditory startle response (ASR) and gap-prepulse inhibition (gap-PPI), tapping temporal processing, were conducted to determine whether EcoHIV infection produced neurocognitive impairments resembling those observed in HIV-1 seropositive individuals.

###### Apparatus

The startle chambers utilized to conduct habituation, cross-modal PPI, and gap-PPI have been previously reported^44^. In brief, a startle platform (SR-Lab Startle Reflex System, San Diego Instruments, Inc., San Diego, CA) was enclosed within an isolation cabinet (external dimensions: 10 cm-thick, double-walled, 81 × 81 × 116-cm) (Industrial Acoustic Company, INC., Bronx, NY), to provide sound attenuation (30 db(A)) relative to the external environment. A SR-Lab system high-frequency loudspeaker (model#40-1278B, Radio Shack, Fort Worth, TX), mounted 30 cm above the Plexiglas animal test cylinder, was used to present continuous background noise (22dB(A)), auditory prepulse stimuli (85 db(A), duration: 20 msec) and the startle stimulus (100 db(A), duration: 20 msec). A 22 lux white LED light, affixed on the wall in front of the test cylinder, was utilized to present visual prepulse stimuli (duration: 20 msec). A piezoelectric accelerometer, attached to the bottom of the Plexiglas animal test cylinder, converted the deflection of the test cylinder, resulting from animal’s response to the startle stimulus, into analog signals. Response signals were digitized (12 bit A to D, recorded at a rate of 2000 samples/sec) and saved to a hard disk. Two individual startle apparatuses were used throughout the duration of experimentation.

###### Procedure

Temporal processing was assessed using cross-modal PPI and auditory gap-PPI in EcoHIV and control animals approximately two weeks after stereotaxic injections.

Following a single session of habituation (methodology reported in Moran et al., 2013), cross-modal PPI and auditory gap-PPI were conducted in a sequential manner. Each session began with a 5-min acclimation period under continuous 70 dB(A) white background noise followed by 6 pulse-only auditory startle response (ASR) trials with a fixed 10 msec intertrial interval (ITI). For cross-modal PPI, a total of 72 trials were presented, including an equal number of auditory and visual prepulse trials. The prestimulus modality (i.e., auditory or visual) was arranged using an ABBA counterbalanced order of presentation. For auditory gap-PPI, a 20-msec gap in background noise served as a discrete prestimulus during 36 testing trials. Trials for both cross-modal PPI and auditory gap-PPI has interstimulus intervals (ISIs) of 0, 30, 50, 100, 200, or 4000 msec, which were presented in 6-trial blocks interdigitated using a Latin Square experimental design. ISIs of 0 and 4000 msec served as control trials to provide a reference ASR within the testing session. A variable (15-25 seconds) ITI was employed for all testing trials. Mean peak ASR amplitude values were collected for analysis.

##### Neurocognitive Assessments: Locomotor Activity

###### Apparatus

Square (40 × 40 cm) activity monitors (Hamilton Kinder, San Diego Instruments, San Diego, CA) were converted into round (~40 cm diameter) compartments using perspex inserts. Interruptions of infrared photocells (32 emitter/detector pairs) were utilized to detect free movement.

###### Procedure

Locomotor activity was conducted on three consecutive days approximately seven weeks after stereotaxic injections of either EcoHIV or saline using a sixty-minute test session. Test sessions were conducted between 7:00 and 12:00 h (EST) in an isolated room under dim lighting conditions (<10 lux).

##### RNAscope in situ Hybridization

Viral infection and replication were verified by labeling HIV-1 mRNA and HIV-1 DNA using the RNAscope *in situ* hybridization methods detailed above. NogoA signaling in EcoHIV (*n*=4, Female, *n*=1, Male, *n*=3) and control (*n*=3, Female, *n*=1, Male, *n*=2) animals was also examined using RNAscope *in situ* hybridization.

##### Neuroanatomical Assessments: Synaptic Dysfunction

Transcardial perfusion, using methodology adapted from Roscoe et al. (2014), was conducted after rats were deeply anesthetized using sevoflurane (Abbot Laboratories, North Chicago, IL) Following perfusion, a ballistic labeling technique, originally described by (Seabold *et al*., 2010) was used to visualize pyramidal neurons from layers II-III of the medial prefrontal cortex (3.7 mm to 2.2 mm anterior to Bregma)^45^. Methodological details on the preparation of ballistic cartridges, Tefzel tubing, and DiOlistic labeling are available in Li et al^46^.

Z-stack images were obtained on three pyramidal neurons and three MSNs from each animal (EcoHIV: *n*=10, Female, *n*=3; Male, *n*=7; Control: *n*=10, Female, *n*=3; Male, *n*=7) using a Nikon TE-2000E confocal microscope and Nikon’s EZ-C1 software (version 3.81b). Based on the selection criteria (i.e., continuous dendritic staining, low background/dye clusters, and minimal diffusion of the DiI dye into the extracellular space), one pyramidal neuron and one MSN from each animal was chosen for analysis. Following selection, Neurolucida 360 (MicroBrightfield, Williston, VT), a sophisticated neuronal reconstruction software, was utilized to examine neuronal morphology and dendritic spine morphology. Specifically, dendritic spine morphology, was assessed using two parameters, including head diameter (μm) and neck diameter (μm).

##### Neuroanatomical Assessments: Signaling Pathways

Key elements of the Nogo-A signaling pathway, which is involved in synaptic plasticity in the central nervous systems^47^, were examined using tissue fluorescence immunostaining techniques (methodology detailed above; EcoHIV: *n*=6, Female, *n*=2; Male, *n*=4; Control: *n*=6, Female, *n*=2, Male, *n*=4). Specifically, brain sections were incubated overnight at 4°C with rabbit anti-PirB (E-AB-15732, Elabscience, Houston, TX, USA), which plays an inhibitory role in axonal regeneration and synaptic plasticity following CNS injury (Atwal et al., 2008), mouse anti-NgR3 (sc-515400, Santa Cruz Biotech, Dallas, TX, USA), whose function is poorly characterized, or mouse anti-Rho A (sc-418, Santa Cruz Biotech, Dallas, TX, USA), whose inhibition is critical to spine maintenance (for review, Lai & Ip, 2013). Then, brain sections were incubated with goat anti-rabbit Alexa Fluor 594 (A11012, Invitrogen, Carlsbad, CA, USA), or donkey anti-mouse Alexa Fluor 594 (A21203, Invitrogen, Carlsbad, USA).

##### Neuroanatomical Assessments: Neuroinflammatory Markers

The total RNA was isolated from 20 mg of brain tissue (EcoHIV: *n*=4, Female, *n*=1 Male, *n*=3; Control: *n*=4, Female, *n*=1, Male, *n*=3) using the RNeasy Mini kit (Cat. No. 74104, QIAGEN, Hilden, German) and manufacturer’s protocol. One μg of total RNA from each sample was converted into cDNA by Cloned AMV first-strand cDNA synthesis kit (Invitrogen, Carlsbad, CA, USA) for real-time PCR. The 20 μL first-strand cDNA synthesis reaction mixture contained 1 μL random-hexamer primer, 1 μg total RNA sample, 2 μL of 10 mM dNTP mix, 5x cDNA synthesis buffer, 1 μL of 0.1 M DTT, 1 μL RNaseOUT, 1 μL cloned AMV RT and 1 μL DEPC-treated water. The following conditions were used: 65 °C for 5 min, 25 °C for 10 min, 50 °C for 50 min and terminated at 85 °C for 5 min. The cDNA products were used immediately for further analysis. The neuroinflammation related cytokines (TNF-α, IL-1β, IL-6) were quantified using a real-time PCR detection system and SsoAdvanced Universal SYBR Green Supermix kit (BIO-RAD). In brief, the 20 μL reaction mixture contained 10 μL 2x SsoAdvanced universal SYBR Green supermix, 1 μL forward primer and reverse primers (250 nM each), 1 μL template (100 ng) and 7 μL of DEPC-treated water. Reactions were performed with the DNA Engine Opticon 2 system (M J Research, USA) using the following cycling conditions: 30 sec at 95 °C and 40 cycles of 15 sec at 95 °C and 30 sec at 59 °C. The data were analyzed using Intuitive Opticon Monitor TM software. All primers are listed in Supplementary Table S1, and β-Actin was used as an internal control. The relative gene expression of each neuroinflammatory marker (i.e., TNF-α, IL-1β, IL-6) was analyzed using the 2^−ΔΔCt^ method to examine relative changes.

### Statistical Analyses

Analysis of variance (ANOVA) and regression techniques were utilized to statistically analyze the data (SAS/STAT Software 9.4, SAS Institute, Inc., Cary, NC, USA; SPSS Statistics 26, IBM Corp., Somer, NY, USA; GraphPad Software, Inc., La Jolla, CA, USA). Figures were created using GraphPad Prism 5 (GraphPad Software Inc., La Jolla, CA, USA. An alpha value of *p*≤0.05 was considered statistically significant for all analyses.

The regional distribution of HIV-1 mRNA in the HIV-1 Tg rat was analyzed using a repeated-measures ANOVA, whereby brain region (i.e., medial prefrontal cortex (mPFC), nucleus accumbens (NAc) and hippocampus (HIP)) served as the within-subjects factors.

Body weight in EcoHIV animals was also analyzed using a repeated measures ANOVA. Age served as the within-subjects factor, while genotype (i.e., EcoHIV vs. Control) and biological sex (i.e., Male vs. Female) served as between-subjects factors. Regression analyses were conducted to evaluate neurocognitive deficits (i.e., temporal processing, long-term episodic memory) in EcoHIV animals.

Dendritic branching complexity, evaluated using a centrifugal branch ordering method, and dendritic arbor complexity, assessed using the classical Sholl analysis^48^, was analyzed using regression analyses. Furthermore, morphological parameters of dendritic spines, including head diameter and neck diameter, were analyzed using a generalized linear mixed effects model with a Poisson distribution with an unstructured covariance pattern (PROC GLIMMIX; SAS/STAT Software 9.4, SAS Institute, Inc., Cary, NC, USA). The number of dendritic spines within each bin served as the dependent variable. Bin served as a within-subjects factor, while genotype (EcoHIV vs. Control) and biological sex (Male vs. Female) served as between-subjects factors. A univariate ANOVA was conducted to evaluate the role of genotype on the NogoA-NgR3/PirB-RhoA signaling pathway. Biological sex was included as a covariate in the analysis of the Nogo A-NgR3/PirB-RhoA signaling pathway.

## RESULTS

### Experiment 1: HIV-1 Transgenic Rats

#### A restricted and region-specific distribution of HIV-1 mRNA was observed in HIV-1 Tg rats

HIV-1 Tg rats exhibited a restricted, region-specific distribution of HIV-1 mRNA, evidenced by RNAscope *in situ* hybridization. HIV-1 expression and intensity were categorized in across multiple brain regions using an ordinal scale, where 0 reflected no HIV-1 mRNA expression and 7 represented high HIV-1 mRNA expression (Figures 2a-2b). Phase-contrast Nissl staining images show the brain regions where HIV-1 mRNA expression was detected (Figure 2c). The green fluorescence signals observed in confocal images (Figure 2d) revealed the restricted HIV-1 mRNA expression (single dot expression pattern) of each marked region labeled using the RNAscope *in situ* hybridization assay.

**Figure 2.**
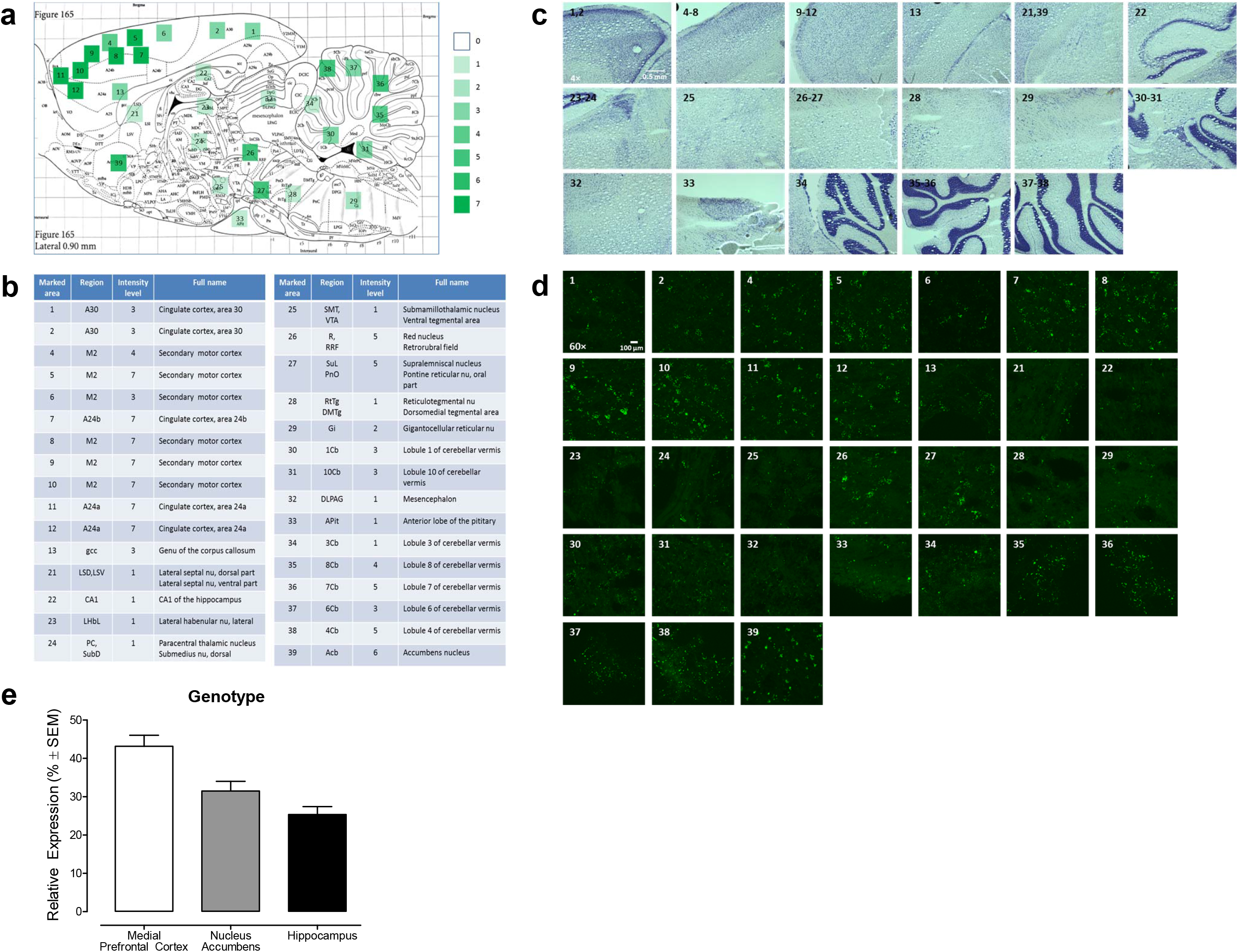
Restricted and region-specific HIV-1 distribution in the HIV-1 Tg rat brain assessed using RNAscope *in situ* hybridization. (a) A sagittal diagram illustrates the brain regions with HIV-1 expression, and intensity, in the HIV-1 Tg rat. (b) All of the expression areas of HIV-1 in HIV-1 Tg rat are listed with the intensity, assessed on a scale from 0 (no HIV-1 expression) to 7 (high HIV-1 expression). The schematic map of sagittal section (lateral 0.90 mm) is cited from *Paxions and Watson’s The Rat brain in Stereotaxic Coordinates* (7th Edition). (c) Representative phase-contrast images (4×) of sagittal brain sections using Nissl staining in brain regions with detected HIV-1 expression are presented. The regions with detectable HIV-1 expression are labeled on the sagittal diagram map above (Figure 1A). (d) Representative confocal images (60×) of HIV-1 expression in different regions of HIV-1 Tg rat. Green fluorescence signal represents the HIV-1 mRNA expression labeled using RNAscope *in situ* hybridization. (e) Statistical analysis of region-specific HIV-1 expression in mPFC, NAc, HIP by gender difference.

The number of green fluorescent dots, representing a single copy of HIV-1 mRNA expression, were counted in three brain regions associated with higher-order cognitive processes, including the medial prefrontal cortex (mPFC)^49^, nucleus accumbens (NAc)^50,51^ and hippocampus (HIP)^52^. The most abundant HIV-1 mRNA expression was observed in the mPFC (Figure 2e). Relative to the mPFC, the nucleus accumbens (NAc) and hippocampus (HIP) exhibited lower levels of HIV-1 mRNA expression. A repeated-measures ANOVA confirmed these observations, revealing a statistically significant main effect of brain region [*F*(2,4)=9.2, p≤0.03, η_p_^2^=0.821] and a main effect of biological sex [*F*(1,2)=178.5, *p*≤0.001, η_p_^2^=0.989]. Notably, other brain regions, including the substantia nigra (SN) and cerebellum also exhibited significant HIV-1 mRNA expression.

#### Cell-type specific HIV-1 mRNA expression was observed in HIV-1 Tg rat

Subsequently, we investigated whether HIV-1 mRNA expression was cell-type specific by conducting RNAscope *in situ* hybridization and immunohistochemical staining for microglia (Iba1) and astrocytes (GFAP) on adjacent brain sections. The majority of green fluorescent dots, reflecting a single copy of HIV-1 mRNA expression, were located inside of microglia, which were identified based on cell morphology (Figure 3a). Unfortunately, due to technological limits, it was not possible to combine the RNAscope *in situ* hybridization assay with immunohistochemical staining in the same sections.

**Figure 3.**
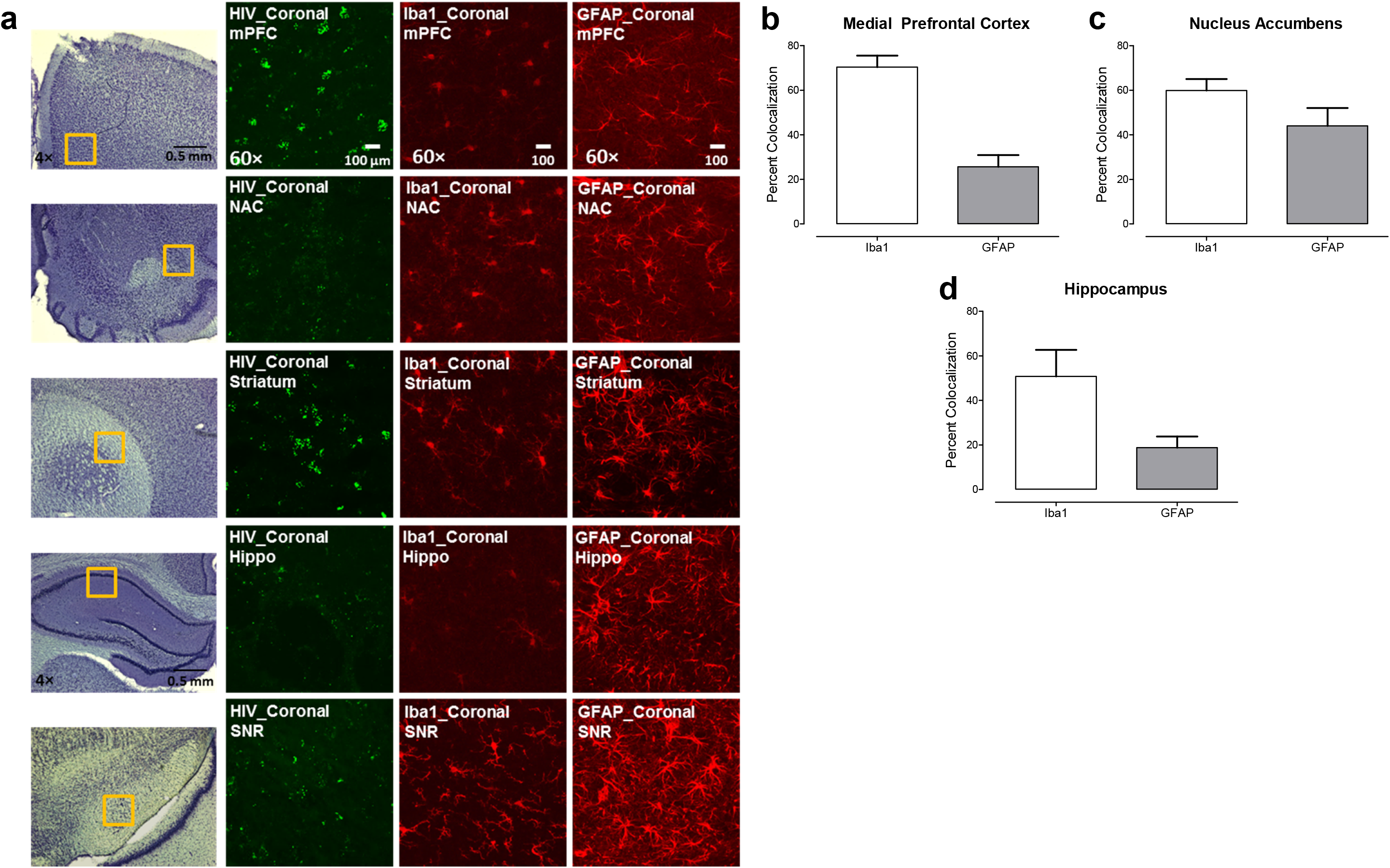
HIV-1 mRNA expression in the mPFC, NAc, HIP and SN of the HIV-1 Tg rat. (a) The yellow frame indicates the selected region, determined via Nissl staining, for confocal imaging. Green fluorescence signal represents HIV-1 mRNA expression. Red fluorescence signals show the cell type (i.e., microglia, astrocytes) based on separate IHC staining for Iba1 and GFAP, respectively. (b-d) Statistical analysis of dual labeling of HIV-1 with each type of cell markers in mPFC, NAc, HIP using RNAscope *in situ* hybridization. The overall percent co-localization and number of HIV-1mRNA copies in different regions and cell types.

In response to these limitations, we utilized the RNAscope dual-labeling assay, affording an opportunity to combine cell-type specific probes (Iba1 and GFAP) with the HIV-1 probe. The percent of co-localization between cell-type specific mRNA expression and HIV-1 mRNA expression were quantified in the mPFC, NAc, and HIP regions in HIV-1 Tg rat brain (Figure 3b–d). Statistical analyses confirmed our observations, revealing that HIV-1 mRNA expression primarily colocalizes with microglia in the mPFC, NAc and HIP. Thus, two complementary techniques, including immunohistochemical staining and the RNAscope *in situ* dual-labeling assay, provide strong, independent evidence for the expression of HIV-1 mRNA in microglia.

#### Purified microglia validated cell-type specific HIV-1 expression

To validate our observations in the HIV-1 Tg rat brain, we isolated and purified microglia from mixed glia cells in HIV-1 Tg rats. Several cell markers specific to microglia (Iba1, MOMA and CD11b) were chosen for immunofluorescence staining to verify the purity of our microglia. Purified microglia (Figure 4a) showed a strong fluorescence signal for all cell markers (Iba1, red; MOMA, green; CD11b, green), indicating reliable purity. Astrocytes (Figure 4A) were labeled with GFAP (green) within the mixed glia culture. Dual-labelling of HIV-1 and Iba1 mRNA using RNAscope *in situ* hybridization (Figure 4b) revealed significant co-localization between HIV-1 and Iba1-positive cells, supporting the role of microglia in brain cells of HIV-1 Tg rats both *in vivo* and *in vitro*.

**Figure 4.**
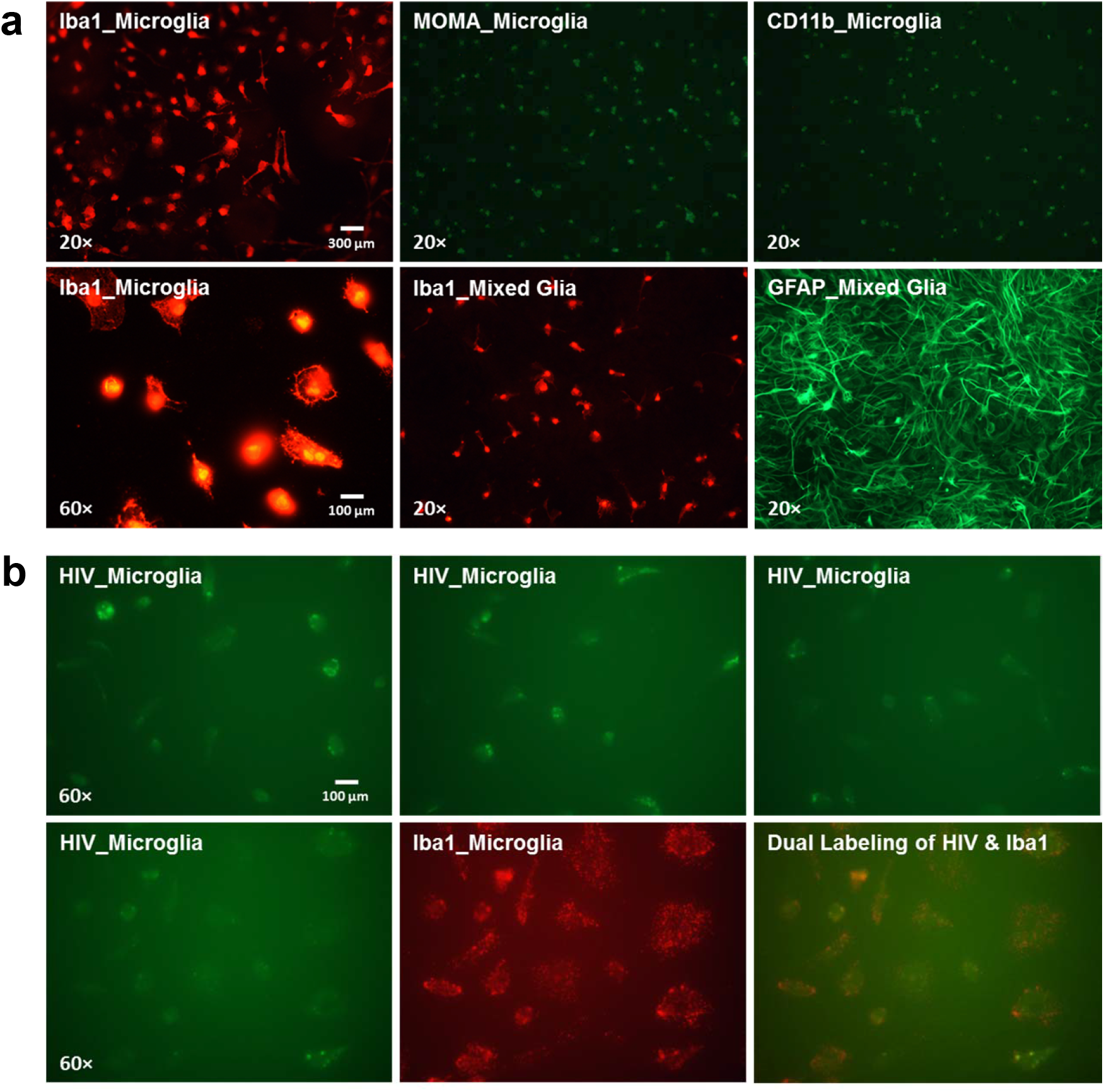
The primary microglia isolation, purification and validation in HIV-1 Tg rat. (a) The representative fluorescence images of the purified microglia cells, which were verified by microglia cell markers: Iba1 (red), MOMA (green), and CD11b (green) at 3 days after purification through IHC staining. The GFAP was used as an indicator of astrocyte in the mixed glia. (b) The representative fluorescence images of the purified microglia cells, which were verified by dual-label of Iba1 and HIV probe through the technique of RNAscope *in situ* hybridization. The three images in the upper panel represented the HIV-1 mRNA expression from three individual experiments by RNAscope *in situ* hybridization. The lower panel showed the co-localization of HIV-1 mRNA with purified microglia cells *in vitro*.

### Experiment 2: EcoHIV Infection Model in Rats

#### Experiment 2.1: In Vitro

##### EcoHIV successfully infects rat primary microglia in vitro

To develop an innovative biological system to model key aspects of HIV-1 infection, we constructed a lentivirus of EcoHIV-EGFP to be applied to healthy rat brain cells *in vitro* and *in vivo*. Figure 5a shows the construction of EcoHIV-EGFP on a backbone of clade B NL4-3 with the replacement of the gp120 gene (the coding region of HIV-1 surface envelope glycoprotein) by ecotropic MLV gp80 gene for entry into cells through CAT-1. The plasmid DNA of EcoHIV-EGFP was transfected into 293FT cells to package the lentivirus (Figure 5b). The conditional medium which included the EcoHIV-EGFP lentivirus was collected and co-cultured with primary microglia isolated from F344/N rat for 24 hours (Figure 5c). EcoHIV-EGFP was well expressed in the microglia (i.e., Iba1 positive cells validated by Iba1 immunostaining), indicating rat primary microglia can be infected by EcoHIV *in vitro*.

**Figure 5.**
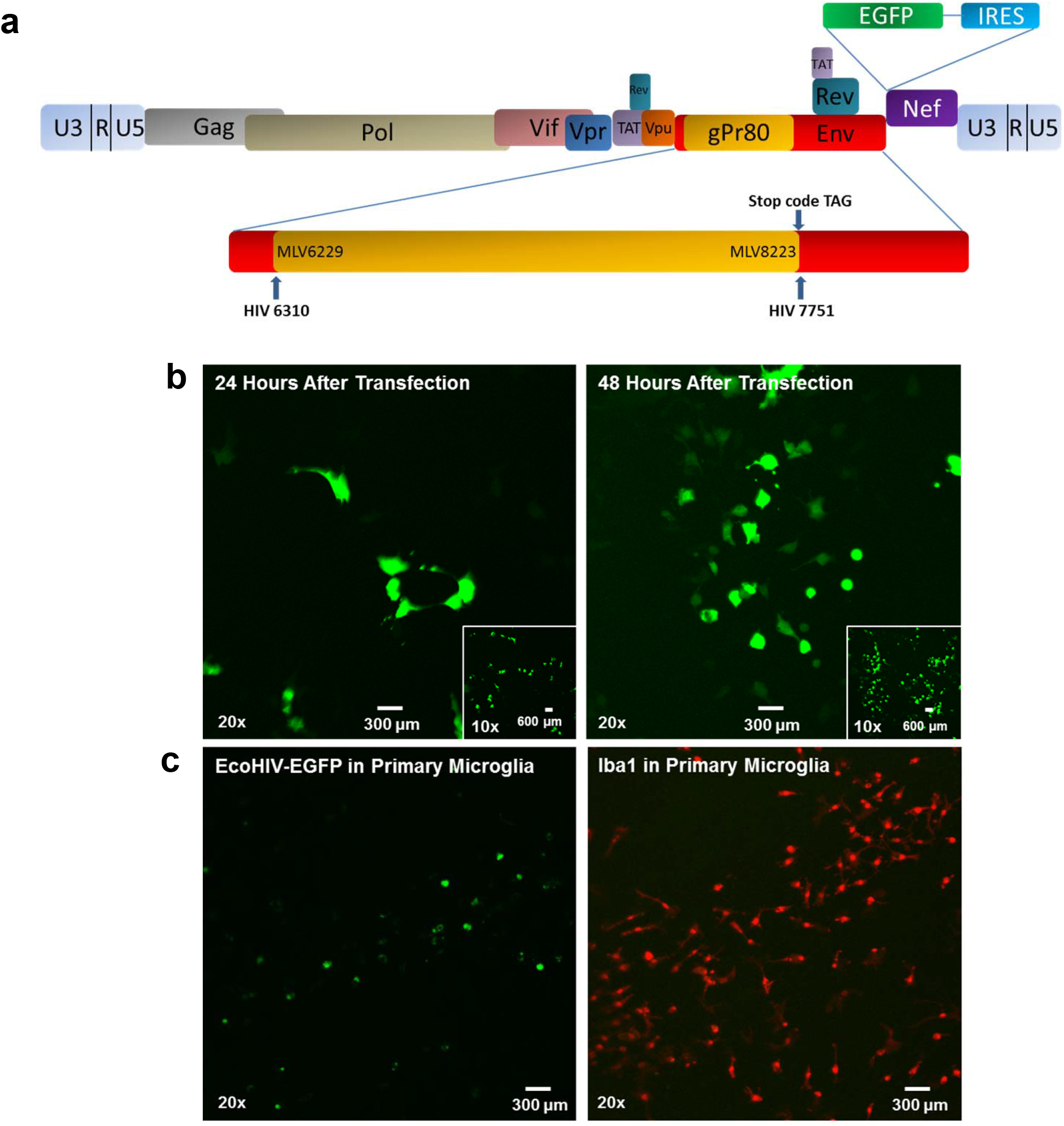
Structure and replication of EcoHIV. (a) Schematic map of EcoHIV-NL4-3-EGFP. The HIV-1 env was replaced by the coding region of the MLV ecotropic envelope gp80 with stop codon. (b) The EcoHIV-EGFP plasmid was transfected into 293FT cell for virus packaging. (c) 24 hours after viral infection, rat primary microglia was immuno- stained with Iba1 antibody.

#### Experiment 2.2: In Vivo, 7 Days Post-Infection

##### Significant infection was observed in microglia seven days after stereotaxic injection of EcoHIV into the rat brain

The conditional medium, including the lentivirus of EcoHIV-EGFP from the infected 293FT cells, was concentrated, tittered, and stereotaxically injected into the cortex of F344/N rats (*n*=6; Female, *n*=2, Male, *n*=4). Seven days after infection, animals were sacrificed and images were taken from coronal brain slices spanning Bregma 5.64 mm to Bregma −4.68 mm revealing significant EcoHIV-EGFP expression throughout the brain (Figure 6a, and Supplementary Figure S1). In both the cortex and the hippocampal dentate gyrus, Iba1 immunostaining co-localized with EcoHIV-EGFP fluorescence signals, providing strong evidence that microglia are the major cell type expressing EcoHIV in the brain (Figure 6b-c). Additionally, retro-orbital injections of EcoHIV-EGFP into F344/N rats (*n*=4; Female, *n*=2, Male, *n*=2) also elicited high expression of EcoHIV-EGFP in both the cortex and hippocampal dentate gyrus after seven days of infection (See Supplementary Figure S2).

**Figure 6.**
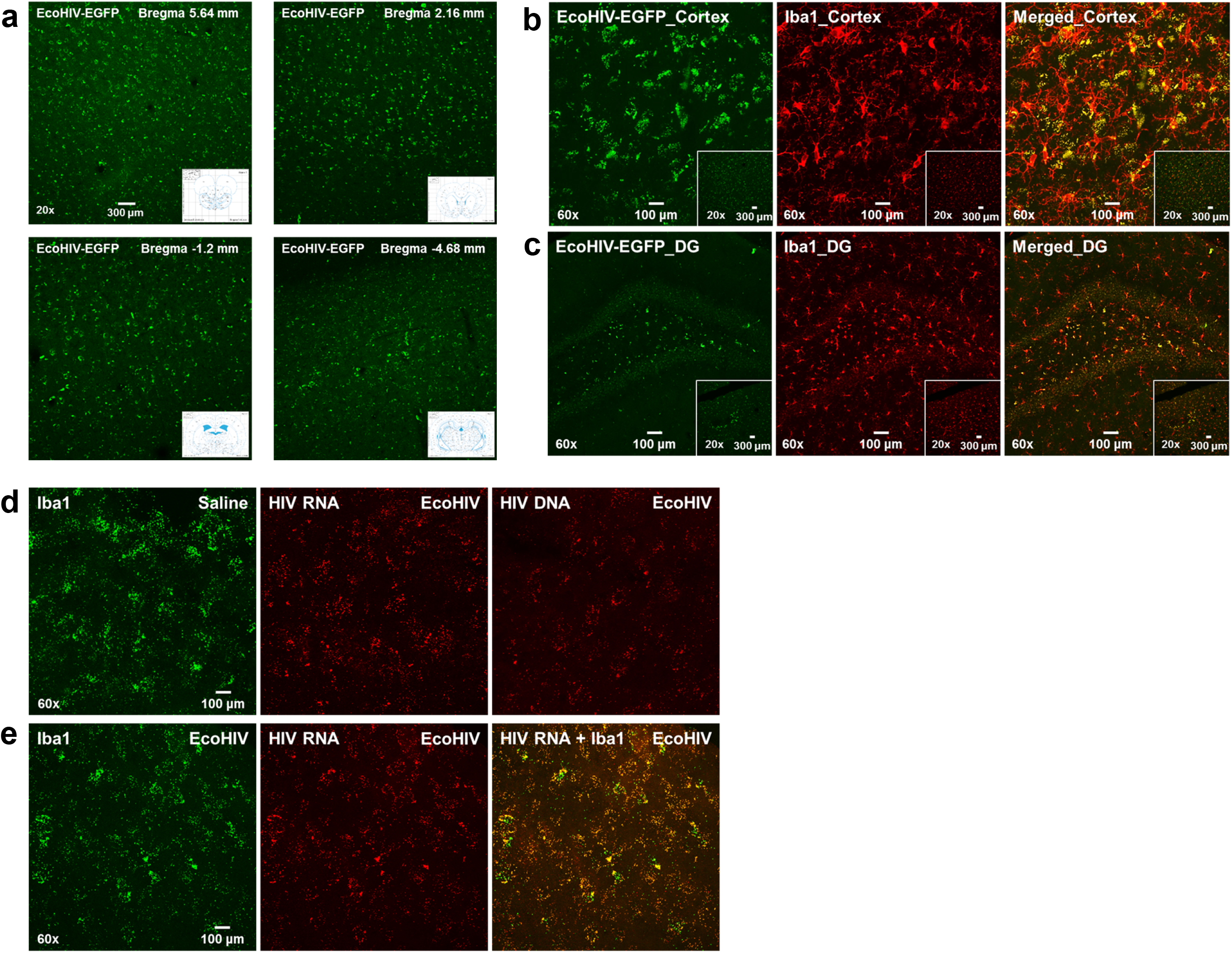
The EcoHIV-EGFP infected cells distributed in rat brain. (a) Four representative confocal images (from different coronal section to bregma: 5.60 mm, 2.16 mm, −1.2 mm, −4.68 mm) of RNAscope *in situ* hybridization show viral infected cells in rat brain at 7 days after the stereotaxic injection. (b-c) The representative images of co-localization of Iba1 immunostaining cortex or hippocampal dentate gyrus regions with EcoHIV-EGFP infected cells at 7 days after injection. (d-e) EcoHIV mRNA or DNA, microglia marker Iba1 expression and dual labeling of EcoHIV mRNA with microglia marker, Iba1, in cortex by RNAscope *in situ* hybridization at 8 weeks after injection.

#### Experiment 2.3: In Vivo, 8 Weeks Post-Infection

##### EcoHIV infection forms viral reservoirs in microglia eight weeks after stereotaxic injections of EcoHIV into the rat brain

The persistence of EcoHIV infection in the brain was assessed using RNAscope *in situ* hybridization to determine 1) if both EcoHIV mRNA and DNA are expressed and 2) the cell type harboring HIV-1 DNA in brain. High levels of HIV-1 mRNA and DNA were observed in the prefrontal cortex, as well as other brain regions (Figure 6d–e). Furthermore, dual-labeling of brain tissue from EcoHIV-infected rats revealed co-localization of HIV mRNA with microglia (i.e., Iba1 positive cells). Most critically, microglia serve as a viral reservoir for HIV-1, evidenced by their ability to harbor EcoHIV DNA in brain tissue (Figure 6d). Collectively, results validate observations in the HIV-1 Tg rat, providing strong evidence for the role of microglia in HIV infection.

##### EcoHIV rats, independent of biological sex, exhibited prominent neurocognitive impairments in temporal processing and long-term episodic memory

The functional health of EcoHIV and control animals was evidenced by using body weight as an assessment of somatic growth. (Figure 7a–b). Both EcoHIV and control animals gained weight throughout the duration of experimentation (Main Effect: Week, [*F*(5, 80)=287.9, *p*≤0.001, η_p_^2^=0.947]). The rate of growth was dependent upon the factor of biological sex (Week × Sex Interaction, [*F*(5, 80)=108.3, *p*≤0.001, η_p_^2^=0.871]). There was no evidence, however, for any effect of EcoHIV infection on somatic growth (*p*>0.05).

**Figure 7.**
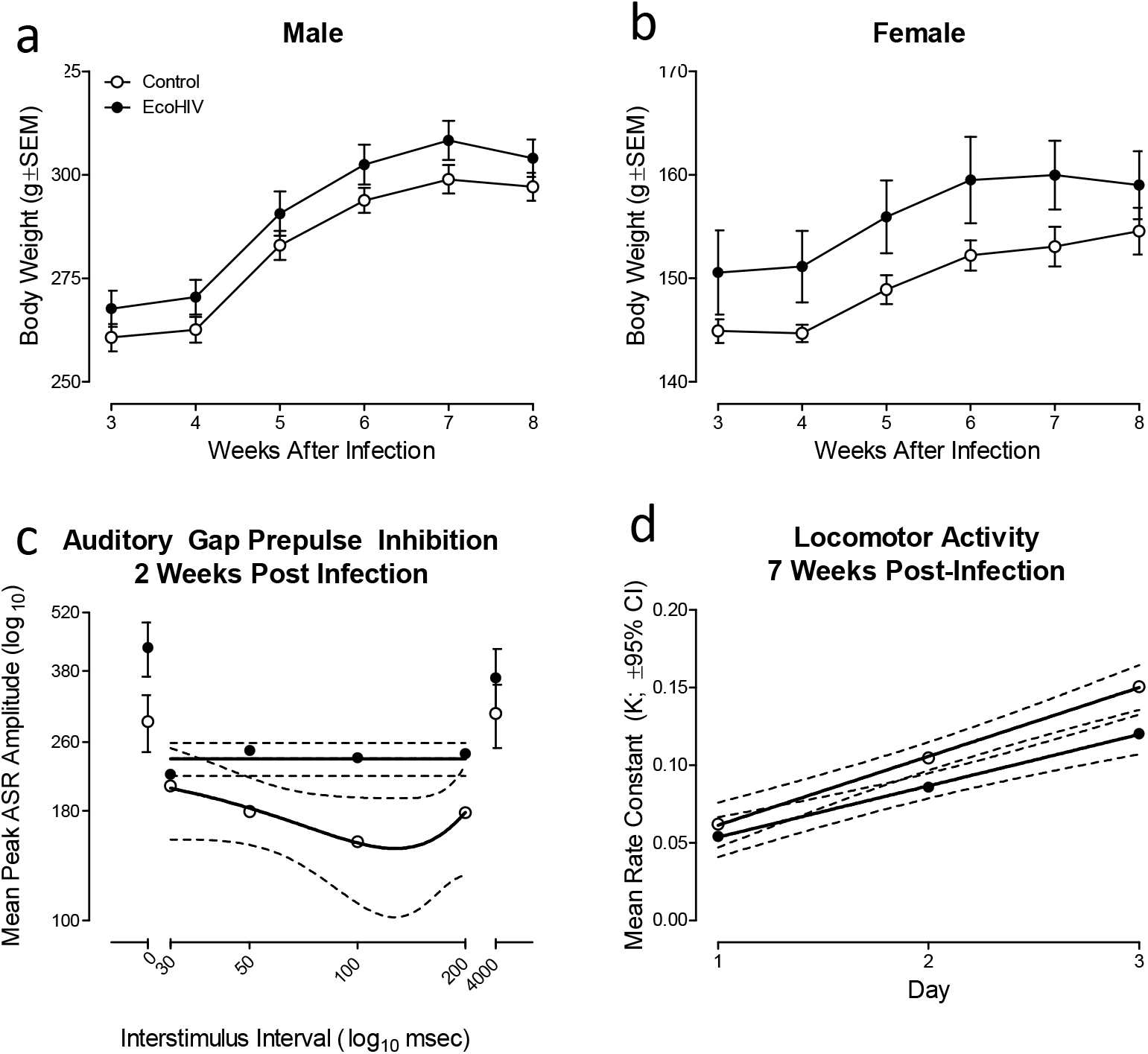
EcoHIV infection induced prominent neurocognitive deficits in temporal processing and long-term episodic memory. (a-b) EcoHIV and control animals, independent of biological sex, exhibited steady growth throughout the duration of the experiment. (c) Auditory gap prepulse inhibition was conducted two weeks after stereotaxic injections of either EcoHIV or saline. EcoHIV infection induced prominent alterations in temporal processing evidenced by the relative insensitivity to the manipulation of interstimulus interval relative to control rats. (d) Three consecutive test sessions in locomotor activity were conducted seven weeks after stereotaxic injections. Alterations in the mean rate constant in EcoHIV rats supports prominent deficits in long-term episodic memory relative to control rats.

Temporal processing, a potential neurobehavioral mechanism underlying alterations in higher-order cognitive functions^32^, was assessed using auditory gap-PPI. EcoHIV animals exhibited a relative insensitivity to the manipulation of ISI relative to control animals (Figure 7c). Regression analyses confirmed these observations, evidenced by differences in the functional relationship describing the inhibition curve during the gap-PPI testing trials (i.e., 30-200 msec ISI). Specifically, a second-order polynomial provided a well-described fit for control animals (*R*^2^≥0.98), with maximal inhibition observed at the 100 msec ISI. A horizontal line, however, afforded the best fit for EcoHIV animals, supporting an insensitivity to the manipulation of ISI. Results support, therefore, a prominent alteration in temporal processing in EcoHIV animals; an alteration that occurs early during the course of viral protein exposure.

Long-term episodic memory, which is commonly altered in HIV-1 seropositive individuals^53,54^, can be assessed using measures of novelty and habituation^55,56,57^. EcoHIV animals exhibited a prominent alteration in long-term episodic memory, evidenced by a differential development of intrasession habituation, relative to control animals. During each locomotor activity test session, a one-phase decay afforded a well-described fit for the mean number of photocell interruptions during locomotor habituation trials across 5-minute trial blocks in both EcoHIV and control animals (*R*^2^s≥0.96). The rate of intrasession habituation (i.e., *K*) during each test session, however, was dependent upon genotype (Figure 7f). Specifically, a first-order polynomial afforded the best fit for the rate of intrasession habituation in both EcoHIV and control animals (*R*^2^s≥0.99). Statistically significant differences in the slope (i.e., β_1_), however, were observed (*F*(1,2)=91.0, *p*≤0.01) supporting a slower development of intrasession habituation in EcoHIV animals. Thus, EcoHIV animals exhibited profound NCI, characterized by alterations in temporal processing and long-term episodic memory.

#### Prominent alterations in neuronal morphology and synaptic function were observed in EcoHIV animals

A ballistic labeling technique and sophisticated neuronal reconstruction software were utilized to examine dendritic spine and neuronal morphology in pyramidal neurons from layers II-III of the mPFC; a brain region associated with higher-order cognitive processes.

EcoHIV infection induced prominent alterations in the morphological parameters (i.e., head diameter, neck diameter) of dendritic spines in pyramidal neurons (Figure 8a-8b). Specifically, EcoHIV infected animals exhibited a population shift towards dendritic spines with increased head diameter (Treatment × Bin Interaction, *F*(12,96)=1.9, *p*≤0.05) and increased neck diameter (Treatment x Bin Interaction, Pyramidal Neurons: *F*(10,80)=2.0, *p*≤0.05) relative to control animals; morphological parameters supporting a population shift towards a ‘stubby’ phenotype. Pyramidal neuronal morphology was also altered by EcoHIV infection, evidenced by profound changes in dendritic branching complexity (Figure 8c) and dendritic arbor complexity (Figure 8D). Dendritic branching complexity was examined using a centrifugal branch ordering method, which assigned each dendrite with a branch order by counting the number of segments traversed. EcoHIV infection increased the frequency of higher-order dendritic branches relative to control animals (Regression Fit: Sigmoidal Dose-Response, *R*^2^s≥0.99; Treatment Differences in Fit: *F*(4,22)=34.7, *p*≤0.001). Furthermore, dendritic arbor complexity was assessed using the classical Sholl analysis^48^. Increased dendritic arbor complexity was observed proximal to the soma in EcoHIV infected animals relative to controls (Regression Fit: Third Order Polynomial, *R*^2^s≥0.86; Treatment Differences in Fit: *F*(4,36)=7.3, *p*≤0.001). In combination, results suggest that infection with EcoHIV may have interfered with synaptogenesis, a process of dendritic and synaptic pruning that occurs in the prefrontal cortex during adolescence and young adulthood^58,59^. Furthermore, it is notable that the factor of biological sex likely interacts with EcoHIV infection, evidenced by a statistically significant treatment x sex x bin interaction (*p*≤0.05) for both head diameter and neck diameter. Given these observations, there is a critical need to conduct additional studies with increased sample size to more fully elucidate the role of biological sex.

**Figure 8.**
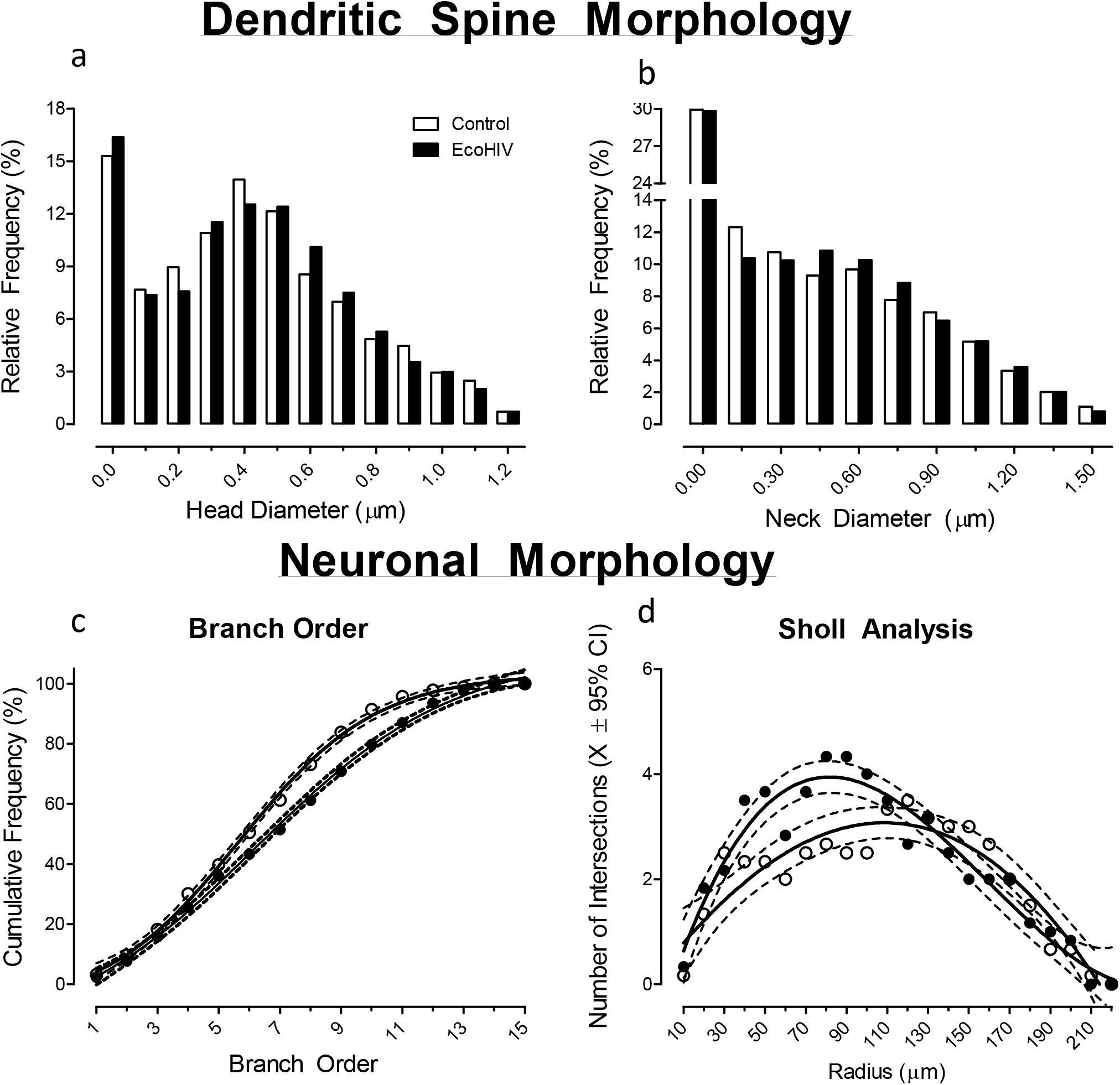
EcoHIV infected animals displayed prominent alterations in dendritic spine and neuronal morphology in pyramidal neurons from layers II-III of the medial prefrontal cortex. (a-b) Rats infected with EcoHIV exhibited a profound morphological shift with an increased relative frequency of dendritic spines with increased head diameter and neck diameter; a morphological shift consistent with a ‘stubby’ dendritic spine phenotype. (c-d) EcoHIV animals exhibited prominent alterations in dendritic branching and neuronal arbor complexity, evidenced by alterations in dendritic branch order and Sholl analysis, respectively.

#### EcoHIV activated the NogoA-NgR3/PirB-RhoA signal pathways, which may mechanistically underlie the prominent alterations in neuronal morphology and synaptic function

Activation of the NogoA-NgR3/PirB-RhoA signaling pathway is involved in the inhibition of axon growth and destabilization of synapses^60^. Eight weeks after EcoHIV infection, the NogoA-mediated signaling pathway was activated, evidenced by the upregulation of NogoA (*F*(1,6)=9.4, *p*≤0.04, η_p_^2^=0.702), NgR3 (*F*(1,12)=16.3, *p*≤0.003, η_p_^2^=0.645), and RhoA (*F*(1,12)=8.1, *p*≤0.02, η_p_^2^=0.475) expression relative to control animals (Figure 9a-d). Alterations in the regulation of PirB were not statistically significant (*p*>0.05). Given the well-recognized role of the NogoA-NgR3/PirB-RhoA signaling pathway in synaptic function, future studies should investigate whether this pathway mechanistically underlies the prominent alterations in neuronal morphology and synaptic function following EcoHIV infection.

**Figure 9.**
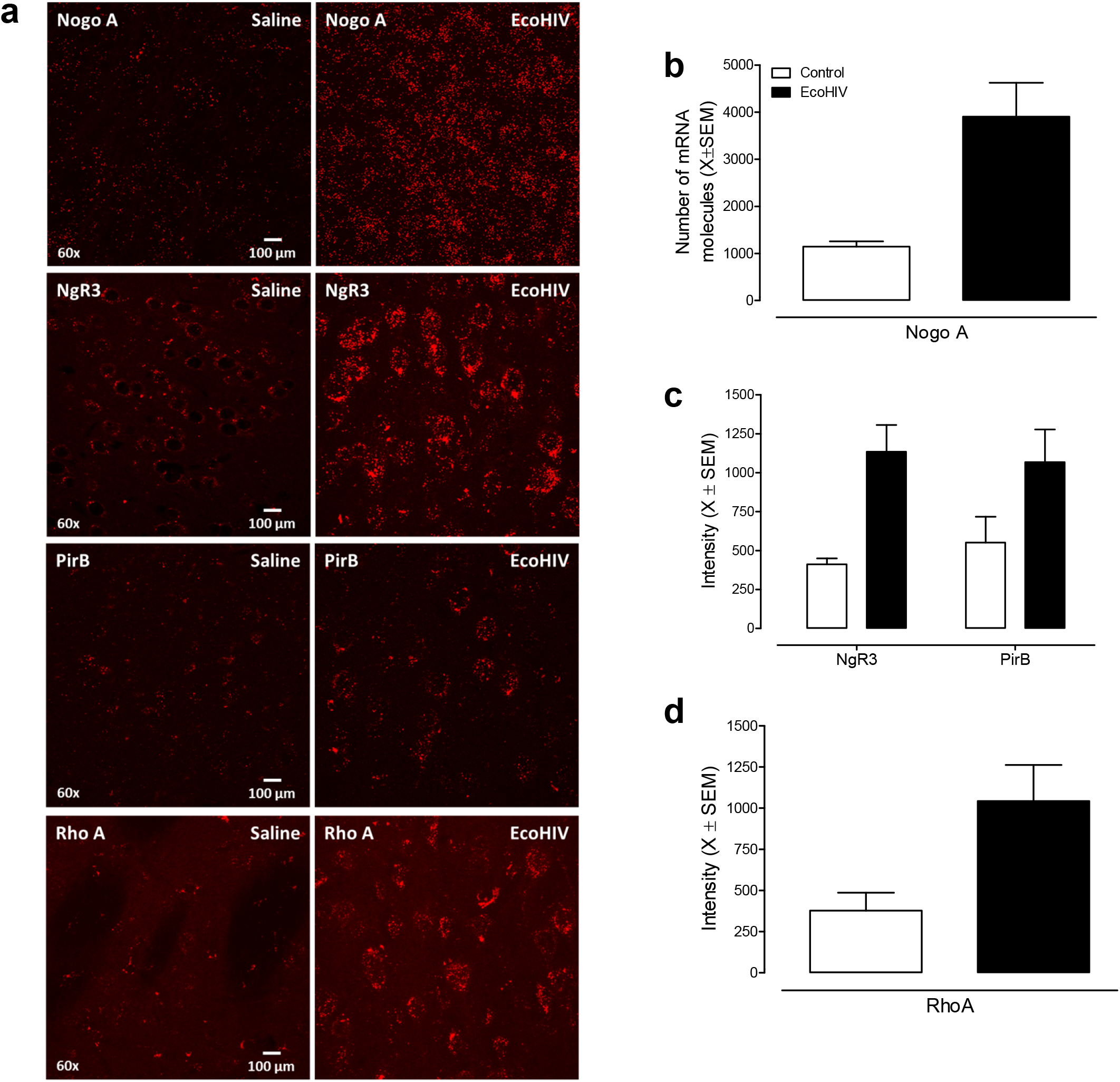
EcoHIV infection activated the NogoA-NgR3/PirB-RhoA signal pathways. (a) Eight weeks after EcoHIV infection, NogoA mRNA was detected by RNAscope *in situ* hybridization in the cortex, NgR3, PirB, and RhoA expression used immunofluorescence staining of EcoHIV infected and control rat. (b-d) Overall, EcoHIV infected rats showed a significant increase of NogoA, NgR3, and RhoA expression in brain compared to control group.

#### A significant difference in neuro-inflammation in brain between EcoHIV infected and control rats

In addition to synaptic dysfunction, neuroinflammation has been implicated as another mechanism underlying HAND. To further establish the utility of the EcoHIV rat as a valid biological system to model HIV infection, four putative neuroinflammatory markers (i.e., NF-κB, TNF-α, IL-6, and IL-1β) were measured in the brain and quantified using the 2^−ΔΔCt^ method. A 1.57-fold increase in NF-κB was detected in the brain tissue of EcoHIV infected animals eight weeks after viral infusion. Furthermore, increased expression levels of TNFα (1.88 fold), IL-1β (1.73 fold), and IL-6 (0.88 fold) in brain were also observed in EcoHIV infected animals (Figure 10a). Inflammatory markers were also examined in spleen (Figure 10b), a key lymphoid organ in the body. EcoHIV animals exhibited increased expression of all four putative neuroinflammatory markers (NF-κB: 2.7 fold; TNF-α: 1.41 fold; IL-6: 2.94 fold; IL-1β: 6.7 fold), relative to the control group. Overall, results support that viral infusion of EcoHIV induced significant neuroinflammation, evidenced by a significant increase in the expression of all four neuroinflammatory genes in both brain and spleen tissue.

**Figure 10.**
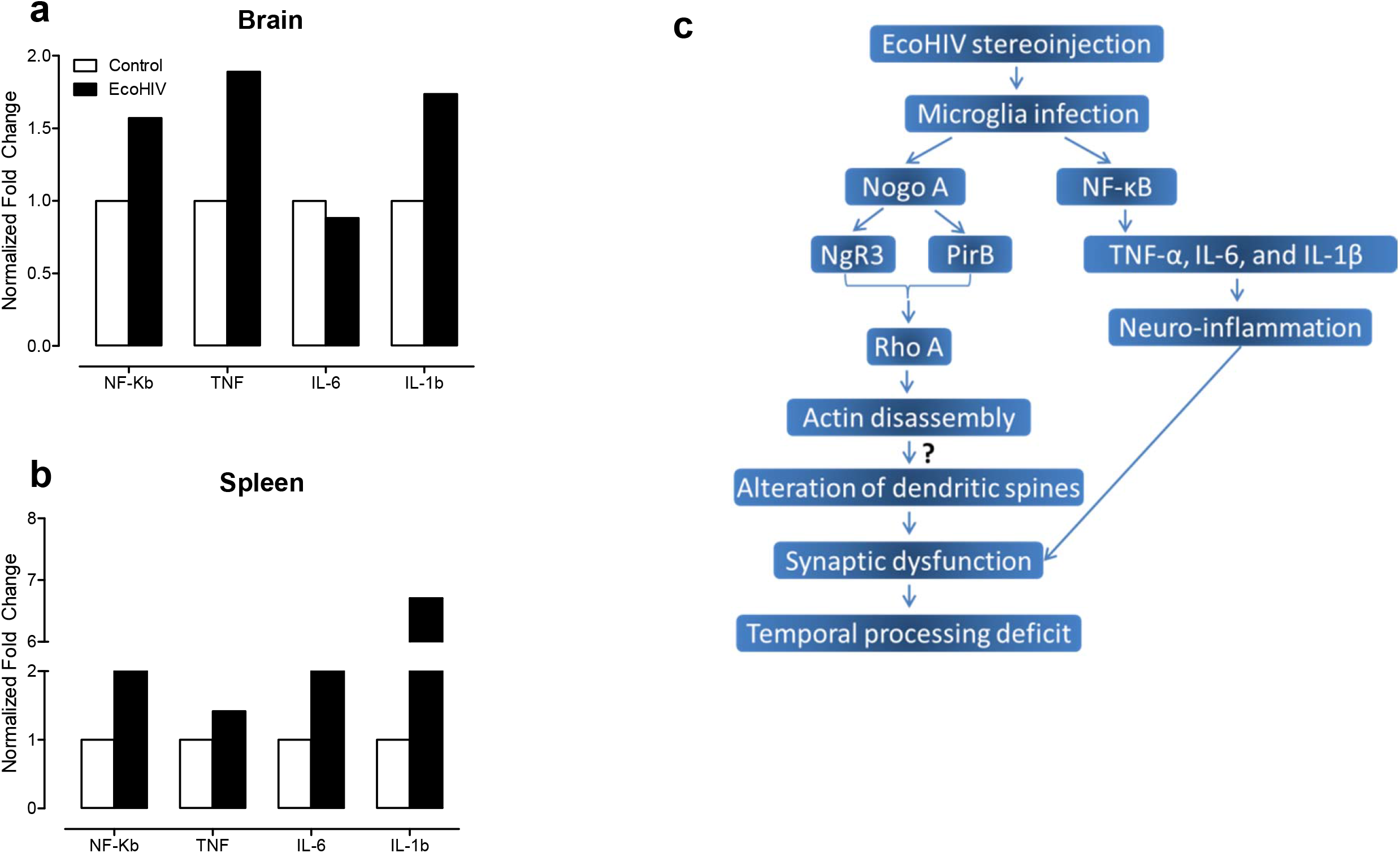
Neuroinflammation in brain between EcoHIV infected and control rats. Eight weeks after EcoHIV infection, four neuroinflammatory markers, including NF-κB, TNF-α, IL-6, IL-1β, were assessed in the cortex (a) and spleen (b) of EcoHIV infected and control animals. Overall, EcoHIV infected rats displayed a significant difference in neuro-inflammation in brain and spleen compared to control group. Data are presented as normalized fold changes. (c) A schematic view of NogoA-NgR3/PirB-RhoA signal pathways during the synaptic dysfunction. After EcoHIV infection, an increase of NogoA could bind to the NgR3 and PirB (which were also increased) following the activation and increase of RhoA, and ultimately leads to the synaptic dysfunction.

## DISCUSSION

Independent of infectivity (i.e., latent virus, as in the HIV-1 Tg rat vs. active infection, as in the EcoHIV rat), microglia are the primary cell-type expressing HIV-1 mRNA and DNA supporting a HIV-1 viral reservoir in the brain; a reservoir which may underlie the persistence of HAND despite cART. In the HIV-1 Tg rat, a restricted, region-specific distribution of HIV-1 mRNA was observed, with the highest level of expression in the mPFC, NAc, SN, cerebellum, and HIP. In EcoHIV rats, an extension of the EcoHIV infection in mice, HIV-1 mRNA was expressed as early as seven days after infection and was localized in microglia. A critical test of the EcoHIV rat across eight weeks revealed the persistence of EcoHIV infection and migration of infection to peripheral organs (i.e., spleen). Neurocognitive and neuroanatomical assessments in EcoHIV rats support the utility of the EcoHIV rat as a biological system to model HAND in the presence of active HIV infection. Collectively, EcoHIV infection in rat replicates observations in the HIV-1 Tg rat, enhancing our understanding of HIV-1 in the brain and offering a novel biological system to model HIV-associated neurocognitive disorders and associated comorbidities (i.e., drug abuse).

With regards to HAND, a common consequence of HIV-1 in the post-cART era, the restricted, region-specific distribution of HIV-1 mRNA in the brain of HIV-1 Tg animals is notable. Across the 39 brain regions that were examined (See Figure 1a–b), the highest levels of mRNA expression were observed in the mPFC, NAc, SN, cerebellum, and HIP; brain regions associated with neurocognitive and neurobehavioral functions commonly altered in HAND. Specifically, the mPFC is associated with multiple neurocognitive functions, including temporal processing^61^, attention^62^ and executive functions^63^; functions that are commonly altered in HIV-1 seropositive individuals^64,^ and the HIV-1 Tg rat^32^. Furthermore, neurocognitive functions related to the hippocampus, including memory, are altered by HIV-1 viral proteins^65,30^. Neurobehavioral alterations associated with the NAc, including motivational dysregulation, are also commonly observed in HIV-1 seropositive individuals^66^ and have been translationally modeled in the HIV-1 Tg rat^67^. Taken together, the highest levels of HIV-1 mRNA expression were observed in brain regions commonly associated with neurocognitive and neurobehavioral functions altered by HIV-1 viral proteins; observations which support the HIV-1 Tg rat as a valid and reliable biological system to model key aspects of HAND in the post-cART era.

Furthermore, brain regions exhibiting high HIV-1 mRNA expression also correspond with neuroanatomical alterations previously observed in the HIV-1 Tg rat. Broadly, the HIV-1 Tg rat exhibits synaptic dysfunction^32,35,44^, neurotransmitter system alterations^68,36^, and neuroinflammation^39^; deficits which have been implicated in the pathogenesis of HAND. More specifically, profound synaptic dysfunction has been observed in both pyramidal neurons from layers II-III of the mPFC^32^ and medium spiny neurons from the NAc^35,44^ evidenced by alterations in dendritic branching complexity and synaptic connectivity in HIV-1 Tg animals relative to controls. Alterations in intrinsic excitability have also been observed in CA1 pyramidal neurons from the HIP^66^. Furthermore, prominent alterations in dopaminergic and serotonergic function have been observed in both the mPFC and NAc^36^. Considering the relatively high expression of HIV-1 mRNA observed in the mPFC, NAc and HIP, even when infection is latent (e.g., as in the HIV-1 Tg rat), the presence and persistence of viral reservoirs may lead to synaptic dysfunction and neurotransmitter system alterations.

Most critically, the utilization of dual-labeling revealed prominent co-localization of HIV-1 mRNA and microglia *in vivo* and *in vitro* supporting microglia as one of the major cell types in which HIV-1 is expressed. Our data indicate that more than 50% of Iba1- positive cells (i.e., microglia) harbor HIV-1 mRNA in the tissues of HIV-1 Tg rats (Figures 3b-d) and *in vitro* purified primary microglia. Additionally, previous studies have reported prominent alterations in both the number^33^ and morphology^69^ of microglia in the HIV-1 Tg rat. The noninfectious nature of the HIV-1 Tg rat, however, precludes HIV-1 viral replication necessitating the extension of a novel biological system (i.e., EcoHIV) to rats for the assessment of the functional role of microglia in HIV-1 infection. Persistent EcoHIV DNA and mRNA were detected in microglia eight weeks after viral infection supporting microglia as a viral reservoir for HIV-1 during active infection; results which are consistent with previous clinical^17,70^ and preclinical^26,27,28,29^.

Furthermore, the current experimental results demonstrate the utility of the EcoHIV rat as a biological system to model HAND in the presence of active HIV infection. First, EcoHIV-EGFP signal was detected in the rat spleen eight weeks after EcoHIV was infused into the brain. Data suggests, therefore, that EcoHIV infection is persistent for at least eight weeks and migrates from brain tissue to infect peripheral organs important in immune defense (Figure 10c; Supplementary Figure S3). Critically, the presence of EcoHIV infection also had functional effects on spleen tissue, evidenced by significant increases in neuroinflammatory markers. Second, the relative health of EcoHIV rats was evidenced by significant growth, independent of biological sex, throughout the experiment. Third, prominent neurocognitive and neuroanatomical deficits were observed in EcoHIV-infected rats within eight weeks of viral infusion. Specifically, EcoHIV-infected rats exhibited profound deficits in temporal processing, which has been implicated as an elemental dimension of HAND^31,32,57^, as well as long-term episodic memory. Anatomically, EcoHIV-infected rats displayed synaptic dysfunction, which may result from alterations in the NogoA-NgR3/PirB-RhoA signaling pathway, and neuroinflammation; functional alterations which may result from microglial activation. Collectively, these results emphasize the utility of the EcoHIV rat as a biological system that can be utilized to model active HIV infection, as well as HAND.

The prominent neuroanatomical alterations observed in the EcoHIV rat merit further discussion given the critical need to elucidate the pathogenesis of HAND^71^. Evaluation of dendritic spine and neuronal morphology in both pyramidal neurons from layers II-III of the mPFC and medium spiny neurons from the NAc revealed prominent synaptic dysfunction. Specifically, synaptic dysfunction was characterized by a shift towards a ‘stubby’ dendritic spine phenotype (i.e., increased dendritic spine head and neck diameter) and an increased frequency of higher-order branches in EcoHIV-infected rats relative to controls. Activation of the NogoA-NgR3/PirB-RhoA signaling pathway, which is involved in axon regeneration and synaptic destabilization^72^, was evidenced by upregulation throughout the signaling pathway. Critically, the binding of NogoA to NgR3 or PirB leads to the activation of RhoA and phosphorylation of PirB, ultimately resulting in actin disassembly and synapse destabilization^73^ affording a potential mechanism underlying synaptic dysfunction in EcoHIV.

Neuroinflammation has also been purported as a potential mechanism underlying HAND^74^. Specifically, in human tissue HIV-1 infection induces activation of the NLRP3 inflammasome and triggers the release of pro-inflammatory cytokines, including IL-1β^75,76^. Furthermore, it has been well documented that the HIV-1 protein, Tat, can activate the NF-κB signaling pathway, subsequently priming NLRP3 inflammasomes and regulating the secretion of cytokines^77,78,79,80^. EcoHIV-infection induced a prominent change in various neuroinflammatory markers (i.e., IL-1β, IL-6, TNF-α, and NF-κB) in the brain relative to control animals. Furthermore, the persistence and peripheral spread of EcoHIV infection was evidenced by the expression of EcoHIV-EGFP in spleen tissue, an important part of the immune-system, and eight weeks after EcoHIV infusion into brain tissue. Since the spleen is one of the important immunosystemic organs, results support the broad utility of EcoHIV-infection in rats as a biological system to model key aspects of HIV-1 and HAND (Figure 10c, and Supplement Figure S3).

The utilization of multiple neuroanatomical assessments afforded a key opportunity to evaluate two potential pathways by which the activation of microglia during active HIV-1 infection may induce neurocognitive impairments (Figure 10c). First, the NogoA-NgR3/PirB-RhoA signaling pathway was activated in EcoHIV infected animals. The NogoA-NgR3/PirB-RhoA signaling pathway underlies actin assembly and thus aberrant activation of this pathway may lead to synaptic dysfunction. Second, prominent changes in various neuroinflammatory markers in the brain support the presence of neuroinflammation, which may also lead to alterations in dendritic spines inducing synaptic dysfunction. In addition, in EcoHIV-infected rats with activated microglia, neurocognitive impairments, characterized by prominent temporal processing deficits, were observed; impairments which potentially result from dendritic spine injuries of the pyramidal neurons in layer II-III of the rat PFC. Notably, since neurons are not infected during HIV infection, damage to dendritic spines more likely results from other infected cell types (i.e., microglia). However, further studies are needed to more directly address the causal nature of these pathways.

Collectively, results of the present experiments identify the most prominent brain regions and cell type targeted by latent HIV in the HIV-1 Tg rat and by active HIV infection in a novel extension of the EcoHIV biological system, the EcoHIV-infected rat. Under conditions of both latent and active HIV-1 infection, microglia are the key cell type harboring HIV supporting a potential novel target for the development of novel therapeutics. Furthermore, the EcoHIV rat, an innovative extension of the EcoHIV mouse, replicated the HIV viral distribution, neurocognitive impairments, and neuroanatomical impairments described throughout the literature on HIV in clinical and preclinical studies. In conclusion, the current experiments have significant implications for our understanding of viral reservoirs and may afford a biological system to advance research on HIV-1, HAND, and the role of comorbidities (i.e., drug abuse).

## ACKNOWLEDGEMENTS

We appreciate Dr. Potash for generous gifts of the EcoHIV-NL4-3-EGFP lentivirus. This work was supported by National Institutes of Health (NIH) grants HD043680, MH106392, DA013137, DA035175 and NS100624. Partial funding was provided by a NIH T32 training grant in Biomedical-Behavioral science.

## AUTHOR CONTRIBUTIONS

HL performed the stereotaxic injection, RNAscope assay and immnunohistochemistry staining, etc.; and was a major contributor in writing the manuscript. KM assisted with the Prepulse inhibition test and data analysis. JI helped with the stereotaxic injection. CM and RB designed the experiment, and revised the manuscript. All authors read and approved the final manuscript.

## REFERENCES

1. Koenig S, et al. Detection of AIDS virus in macrophages in brain tissue from AIDS patients with encephalopathy. Science 233(4768):1089–93 (1986).

2. Whitney JB, et al. Rapid seeding of the viral reservoir prior to SIV viraemia in rhesus monkeys. Nature 512(7512):74–7 (2014).

3. Williams DW, et al. Monocyte maturation, HIV susceptibility, and transmigration across the blood brain barrier are critical in HIV neuropathogenesis. J. Leukoc. Biol. 91(3):401–15 (2012).

4. Cosenza MA, et al. Human brain parenchymal microglia express CD14 and CD45 and are productively infected by HIV-1 in HIV-1 encephalitis. Brain Pathol. 12(4):442–55 (2002).

5. Arts EJ, et al. Cold Spring Harb Perspect Med. 2(4): a007161 (2012).

6. Gray LR, et al. Is the central nervous system a reservoir of HIV-1? Curr Opin HIV AIDS 9(6):552–8 (2014).

7. Smit TK, et al. Independent evolution of human immunodeficiency virus (HIV) drug resistance mutations in diverse areas of the brain in HIV-infected patients, with and without dementia, on antiretroviral treatment. J. Virol. 78(18):10133–48 (2004).

8. Strain MC, et al. Genetic composition of human immunodeficiency virus type 1 in cerebrospinal fluid and blood without treatment and during failing antiretroviral therapy. J Virol. 79(3):1772–88 (2005).

9. Salemi M, et al. Phylodynamic analysis of human immunodeficiency virus type 1 in distinct brain compartments provides a model for the neuropathogenesis of AIDS. J Virol. 79(17):11343–52 (2005).

10. Lamers SL, et al. HIV-1 nef protein visits B-cells via macrophage nanotubes: a mechanism for AIDS-related lymphoma pathogenesis? Curr HIV Res. 8(8):638–40 (2010).

11. Polazzi E, et al. Microglia and neuroprotection: from in vitro studies to therapeutic applications. Prog. Neurobiol. 92(3):293–315 (2010).

12. Réu P, et al. The Lifespan and Turnover of Microglia in the Human Brain. Cell Rep. 20(4):779–784 (2017).

13. Moy J, et al. Temporal effects of estradiol and diethylstilbestrol on pituitary and plasma prolactin levels in ovariectomized Fischer 344 and Holtzman rats: a comparison of radioimmunoassay and Nb2 lymphoma cell bioassay. Proc Soc Exp Biol Med. 200(4):507–13 (1992).

14. Cenker JJ, et al. Brain microglial cells are highly susceptible to HIV-1 infection and spread. AIDS Res. Hum. Retroviruses. 33:1155–1165 (2017).

15. Wallet C, et al. Microglial Cells: The Main HIV-1 Reservoir in the Brain. Front. Cell. Infect. Microbiol. 9:362 (2019).

16. Ko A, et al. Macrophages but not Astrocytes Harbor HIV DNA in the Brains of HIV-1-Infected Aviremic Individuals on Suppressive Antiretroviral Therapy. J. Neuroimmune. Pharmacol. 14(1):110–119 (2019).

17. Churchill MJ, et al. Transcriptional activity of blood-and cerebrospinal fluid-derived nef/long-terminal repeat sequences isolated from a slow progressor infected with nef-deleted human immunodeficiency virus type 1 (HIV-1) who developed HIV-associated dementia. J Neurovirol. 12(3):219–28 (2006).

18. Wohleb ES. Neuron-Microglia Interactions in Mental Health Disorders: “For Better, and For Worse”. Front. Immunol. 7:544 (2016).

19. Domercq M, et al. Neurotransmitter signaling in the pathophysiology of microglia. Front. Cell Neurosci. 7:49 (2013).

20. Streit WJ, et al. Microglia and neuroinflammation: a pathological perspective. J Neuroinflammation. 1(1):14 (2004).

21. Masliah E, et al. Dendritic injury is a pathological substrate for human immunodeficiency virus-related cognitive disorders. HNRC Group. The HIV Neurobehavioral Research Center. Ann. Neurol. 42(6):963–72 (1997).

22. Gelman BB, et al. The National NeuroAIDS Tissue Consortium brain gene array: two types of HIV-associated neurocognitive impairment. PLoS One. 7(9):e46178 (2012).

23. Vera JH, et al. Neuroinflammation in treated HIV-positive individuals: A TSPO PET study. Neurology. 86(15):1425–1432 (2016).

24. Eisele E, et al. Redefining the viral reservoirs that prevent HIV-1 eradication. Immunity. 37(3):377–88 (2012).

25. Sopper S, et al. The effect of simian immunodeficiency virus infection in vitro and in vivo on the cytokine production of isolated microglia and peripheral macrophages from rhesus monkey. Virology. 220(2):320–9 (1996).

26. Micci L, et al. CD4 depletion in SIV-infected macaques results in macrophage and microglia infection with rapid turnover of infected cells. PLoS Pathog. 10:e1004467 (2014).

27. Avalos CR, et al. Brain Macrophages in Simian Immunodeficiency Virus-Infected, Antiretroviral-Suppressed Macaques: a Functional Latent Reservoir. mBio. 8(4):e01186–17 (2017).

28. Llewellyn GN, et al. HIV-1 infection of microglial cells in a reconstituted humanized mouse model and identification of compounds that selectively reverse HIV latenc. J Neurovirol. 24(2):192–203 (2018).

29. Gu CJ, et al. EcoHIV infection of mice establishes latent viral reservoirs in T cells and active viral reservoirs in macrophages that are sufficient for induction of neurocognitive impairment. PLoS Pathog. 14(6): e1007061 (2018).

30. Vigorito M, et al. Spatial learning and memory in HIV-1 transgenic rats. J Neuroimmune Pharmacol. 2(4):319–28 (2007).

31. Moran LM, Booze RM, Mactutus CF. Time and time again: temporal processing demands implicate perceptual and gating deficits in the HIV-1 transgenic rat. J. Neuroimmune. Pharmacol. 8(4):988–97 (2013).

32. McLaurin KA, et al. Disruption of Timing: NeuroHIV Progression in the Post-cART Era. Sci Rep. 9(1):827 (2019).

33. Repunte-Canonigo V, et al. Gene expression changes consistent with neuroAIDS and impaired working memory in HIV-1 transgenic rats. Mol. Neurodegener. 9:26 (2014).

34. Reid W, et al. An HIV-1 transgenic rat that develops HIV-related pathology and immunologic dysfunction. Proc. Natl. Acad. Sci. USA. 98(16):9271–6 (2001).

35. Roscoe RF, et al. HIV-1 transgenic female rat: Synaptodendritic alterations of medium spiny neurons in the nucleus accumbens. J. Neuroimmune. Pharmacol. 9:642–653 (2014).

36. Denton AR, et al. Selective monoaminergic and histaminergic circuit dysregulation following long-term HIV-1 protein exposure. J. Neurovirol. 25(4):540–550 (2019).

37. Peng J, et al. The HIV-1 transgenic rat as a model for HIV-1 infected individuals on HAART. J. Neuroimmunol. 218(1-2):94–101 (2010).

38. Abbondanzo SJ, et al. HIV-1 transgenic rats display alterations in immunophenotype and cellular responses associated with aging. PLoS One. 9(8):e105256 (2014).

39. Royal W, et al. Immune activation, viral gene product expression and neurotoxicity in the HIV-1 transgenic rat. J Neuroimmunol. 247(1-2):16–24 (2012).

40. Potash MJ, et al. A mouse model for study of systemic HIV-1 infection, antiviral immune responses, and neuroinvasiveness. Proc. Natl. Acad. Sci. USA. 102(10):3760–5 (2005).

41. Geraghty P, et al. HIV infection model of chronic obstructive pulmonary disease in mice. Am. J. Physiol. Lung Cell Mol. Physiol. 312(4):L500–L509 (2017).

42. Li H, et al. Identification of Dopamine D1-Alpha Receptor Within Rodent Nucleus Accumbens by an Innovative RNA In Situ Detection Technology. J Vis Exp. 27(133):57444 (2018).

43. Moussaud S, et al. A new method to isolate microglia from adult mice and culture them for an extended period of time. J. Neurosci. Methods. 187(2):243–53 (2010).

44. McLaurin KA, et al. Evolution of the HIV-1 transgenic rat: utility in assessing the progression of HIV-1-associated neurocognitive disorders. J Neurovirol. 24(2):229–245 (2018).

45. Paxinos G. & Watson C. The rat brain in stereotaxic coordinates. 7th ed. (Elsevier Academic Press; Burlington (2014).

46. Li H, et al. Ballistic Labeling of Pyramidal Neurons in Brain Slices and in Primary Cell Culture. J Vis Exp. (158) (2020).

47. Kempf A, et al. Nogo-A represses anatomical and synaptic plasticity in the central nervous system. Physiology (Bethesda). 28(3):151–63 (2013).

48. Sholl A, et al. Pattern discrimination and the visual cortex. Nature 171(4348):387–8 (1953).

49. Grossmann T. The role of medial prefrontal cortex in early social cognition. Front. Hum. Neurosci. 7:340 (2013).

50. Floresco SB. The nucleus accumbens: an interface between cognition, emotion, and action. Annu Rev. Psychol. 66:25–52 (2015).

51. Li H, et al. Posterior ventral tegmental area-nucleus accumbens shell circuitry modulates response to novelty. PLoS One. 14(3): e0213088 (2019).

52. Olsen RK, et al. The hippocampus supports multiple cognitive processes through relational binding and comparison. Front Hum Neurosci. 6:146 (2012).

53. McLaurin KA, et al. Unraveling individual differences in the HIV-1 transgenic rat: Therapeutic efficacy of methylphenidate. Sci Rep. 8:136 (2018).

54. Morgan EE, et al. HIV-associated episodic memory impairment: evidence of a possible differential deficit in source memory for complex visual stimuli. J. Neuropsychiatry. Clin. Neurosci. 21(2):189–98 (2009).

55. Eacott MJ, et al. Recollection in an episodic-like memory task in the rat. Learn Mem. 12(3):221–3 (2005).

56. Barker JC, et al. Experiences of gamma hydroxybutyrate (GHB) ingestion: a focus group study. J. Psychoactive. Drugs. 39(2):115–29 (2007).

57. Chao A, et al. Malnutrition and Nutritional Support in Alcoholic Liver Disease: a Review. Curr Gastroenterol Rep. 18(12):65 (2016).

58. Huttenlocher PR. Synaptic density in human frontal cortex - developmental changes and effects of aging. Brain Res. 163(2):195–205 (1979).

59. Drzewiecki CM, Willing J, Juraska JM. Synaptic number changes in the medial prefrontal cortex across adolescence in male and female rats: A role for pubertal onset. Synapse 70(9):361–8 (2016).

60. Guo D, et al. Spinal presynaptic inhibition in pain control. Neuroscience. 283:95–106 (2014).

61. Ellenbroek B, et al. Rodent models in neuroscience research: is it a rat race? Dis Model Mech. 9(10):1079–1087 (2016).

62. Kim WJ, et al. Utility of the Montreal Cognitive Assessment (MoCA) and its subset in HIV-associated neurocognitive disorder (HAND) screening. J Psychosom. Res. 80:53–7 (2016).

63. Jin et al. Cognitive and cortical thinning patterns of subjective cognitive decline in patients with and without Parkinson’s disease. Parkinsonism. Relat. Disord. 20(9):999–1003 (2014).

64. Heaton RK, et al. HIV-associated neurocognitive disorders before and during the era of combination antiretroviral therapy: differences in rates, nature, and predictors. J Neurovirol. 17(1):3–16 (2011).

65. Maki PM, et al. Impairments in memory and hippocampal function in HIV-positive vs HIV-negative women: a preliminary study. Neurology 72(19):1661–8 (2009).

66. Kamat R, et al. Implications of apathy and depression for everyday functioning in HIV/AIDS in Brazil. J. Affect. Disord. 150(3):1069–75 (2013).

67. Bertrand L, et al. Targeting the HIV-infected brain to improve ischemic stroke outcome. Nat. Commun. 10(1):2009 (2019).

68. Javadi-Paydar M, et al. Locomotor and reinforcing effects of pentedrone, pentylone and methylone in rats. Neuropharmacology 134(Pt A):57–64 (2018).

69. Sokolova IV, et al. Reduced intrinsic excitability of CA1 pyramidal neurons in human immunodeficiency virus (HIV) transgenic rats. Brain Res. 1724:146431 (2019).

70. Rowson SA, et al. Neuroinflammation and Behavior in HIV-1 Transgenic Rats Exposed to Chronic Adolescent Stress. Front Psychiatry. 7:102 (2016).

71. Cosenza MA, et al. Human brain parenchymal microglia express CD14 and CD45 and are productively infected by HIV-1 in HIV-1 encephalitis. Brain Pathol. 12(4):442–55 (2002).

72. Saylor D, et al. HIV-associated neurocognitive disorder--pathogenesis and prospects for treatment. Nat. Rev. Neurol. 12(4):234–48 (2016).

73. Akbik F, et al. Myelin associated inhibitors: a link between injury-induced and experience-dependent plasticity. Exp. Neurol. 235(1):43–52 (2012).

74. Taylor J, et al. The scaffold protein POSH regulates axon outgrowth. Mol Biol Cell. 19(12):5181–92 (2008).

75. Hong S, et al. Role of the immune system in HIV-associated neuroinflammation and neurocognitive implications. Brain Behav Immun. 45:1–12 (2015).

76. Walsh JG, et al. Rapid inflammasome activation in microglia contributes to brain disease in HIV/AIDS. Retrovirology 11:35 (2014).

77. Maingat FG, et al. Neurosteroid-mediated regulation of brain innate immunity in HIV/AIDS: DHEA-S suppresses neurovirulence. FASEB J. 27:725–737 (2013).

78. Chivero ET, et al. HIV-1 Tat Primes and Activates Microglial NLRP3 Inflammasome-Mediated Neuroinflammation. J Neurosci. 37(13):3599–3609 (2017).

79. Fiume G, et al. Human immunodeficiency virus-1 Tat activates NF-kappaB via physical interaction with IkappaB-alpha and p65. Nucleic Acids Res. 40:3548–3562 (2012).

80. Dandekar DH, et al. HIV-1 Tat directly binds to NFkappaB enhancer sequence: role in viral and cellular gene expression. Nucleic Acids Res. 32:1270–1278 (2004).

